# Biochemical and structural analysis reveals the Neurofibromin (NF1) protein forms a high-affinity dimer

**DOI:** 10.1101/757856

**Authors:** Mukul Sherekar, Sae-Won Han, Rodolfo Ghirlando, Simon Messing, Matthew Drew, Puneet Juneja, Hugh O’Neill, Christopher Stanley, Debsindhu Bhowmik, Arvind Ramanathan, Sriram Subramaniam, Dwight V. Nissley, William Gillette, Frank McCormick, Dominic Esposito

**Affiliations:** NCI RAS Initiative, Cancer Research Technology Program, Frederick National Laboratory for Cancer Research, Frederick, MD, 21702; Helen Diller Comprehensive Cancer Center, University of California San Francisco, San Francisco, CA, 94158; Department of Internal Medicine, Seoul National University Hospital, South Korea; Laboratory of Molecular Biology, NIDDK, NIH, Bethesda, MD, 20892; Robert P. Apkarian Integrated Electron Microscopy Core, Emory University, Atlanta, GA, 30322; Oak Ridge National Laboratory, Oak Ridge, TN, 37830; Argonne National Laboratory, Lemont, IL, 60439; Frederick National Laboratory for Cancer Research, Frederick, MD, 21702; University of British Columbia, Vancouver, British Columbia, V6T1Z3, Canada

**Author notes:** Both authors contributed equally to this manuscript. To whom correspondence should be addressed: Frederick National Laboratory for Cancer Research, PO Box B, Frederick, MD 21702. The abbreviations used are: NF1, Neurofibromatosis Type 1; MAPK, mitogen-activated protein kinase; GAP, GTPase-activating protein; GRD, GAP-related domain; PH, pleckstrin homology; SEC, size exclusion chromatography; SEC-MALS, size exclusion chromatography multi-angle light scattering; SAXS, small-angle X-ray scattering; SANS, small-angle neutron scattering, TEM, transmission electron microscopy; AUC, analytical ultracentrifugation; IP, immunoprecipitation; MBP, maltose-binding protein; TEV, tobacco etch virus.

**Keywords:** NF1, Neurofibromin, RASopathy, GTPase-activating protein, RAS, KRAS, GTPase, MAPK pathway, protein dimerization

## Abstract

Neurofibromin is the protein product of the *NF1* gene which is mutated in the Rasopathy disease Neurofibromatosis Type I. Defects in *NF1* lead to aberrant signaling through the RAS-MAPK pathway due to disruption of the Neurofibromin GTPase-activating function on RAS family small GTPases. Very little is known about the function of the majority of Neurofibromin—to date, biochemical and structural data exist only for the GAP domain and the region containing a Sec-PH motif. To better understand the role of this large protein, we carried out a series of biochemical and biophysical studies which demonstrate that full length Neurofibromin forms a high-affinity dimer. Neurofibromin dimerization also occurs in cells, and likely has biological and clinical implications. Analysis of purified full-length and truncated variants of Neurofibromin by negative stain electron microscopy reveals the overall architecture of the dimer and predicts the potential interactions which contribute to the dimer interface. Structures resembling high-affinity full-length dimers could be reconstituted by mixing N- and C-terminal protein domains *in vitro*. Taken together these data suggest how Neurofibromin dimers might form and be stabilized within the cell.

## INTRODUCTION

Neurofibromatosis type 1, a disease characterized by benign neurofibromas and malignant tumors of the nervous system, is caused by mutations in the 300 kB tumor suppressor gene, *NF1*, that encodes the protein Neurofibromin (1). NF1 is also mutated in various types of cancer, including melanoma (2), leukemia (3), and lung (4). Neurofibromin is a large, multi-domain protein with its best described activity being the down regulation of the MAPK pathway through its GTPase activating protein (GAP) domain (5,6), which stimulates RAS GTPase activity to shut off signaling. Given that there are no mutational hotspots in *NF1* (with over 2,000 mutations reported in the Human Gene Mutation Database) (7) and that the phenotypes reported for mutations are diverse (reviewed in (8)), it is possible that other *NF1* domains have a role in human disease. Thus, increasing the basic knowledge of Neurofibromin structural biology would likely stimulate progress in this field considerably.

Unsurprisingly for such a large multidomain protein, Neurofibromin is reported to interact with several proteins: SPRED family proteins (9), tubulin (10), kinesin-1 (11), protein kinase A (12), protein kinase C (13), caveolin (14), and amyloid precursor protein (15). Accordingly, of the several domains identified in Neurofibromin by homology (cysteine/serine rich domain, tubulin-binding domain, GAP-related domain, Sec-PH domain, focal adhesion kinase-interacting domain), several are known to be protein-protein interaction domains. To date, detailed structural information on Neurofibromin is limited to high-resolution crystal structures of the GAP domain (16), the Sec-PH domain (17,18), and the ternary complex between KRAS4b, the EVH1 domain of SPRED1, and the GRD from NF1 (Wupeng Yan, personal communication). No detailed information exists about the structure or function of the remaining 80% of the protein.

Towards this end, we report here the production of purified full length Neurofibromin and a number of fragments representing various domains of the protein. We provide *in vitro* and cell-based data that demonstrate Neurofibromin exists as a high-affinity dimer. From electron microscopy experiments on full-length Neurofibromin and domains, we propose a possible model for organization of the Neurofibromin dimer and identify regions of importance for dimer formation. Collectively, this work supports the notion that a stable Neurofibromin dimer exists in cells and provides a foundation to elucidate mechanisms underlying the role of Neurofibromin in the diverse collection of human diseases associated with this protein.

## EXPERIMENTAL PROCEDURES

### Cloning of Neurofibromin constructs

DNA constructs were generated for the various fragments and full-length Neurofibromin using Gateway recombinational cloning (19). Entry clones were produced by PCR amplification of portions of the NF1 gene from Addgene construct 70423, which contains a codon optimized version of human NF1 isoform 2 (NM_000267.3) to ensure stability of the plasmid in *E. coli*. 5’ sequences included Gateway *att*B1 sites followed by sequences for protease cleavage by tobacco etch virus (TEV) protease (ENLYFQ/G), while 3’ sequences contained stop codons and Gateway *att*B2 sites. PCR amplicons were cloned into pDonr255 and fully sequence verified. Gateway LR recombination was used to generate final expression clones in baculovirus Destination vectors pDest-635 (pFastBac1 with N-terminal His6 tag) and pDest-636 (pFastBac1 with N-terminal His6-MBP tag), or E. coli Destination vector pDest-566 (N-terminal His6-MBP tag, Addgene #11517). Final baculovirus expression clones were transformed into DH10Bac cells (ThermoFisher), and bacmid clones were verified by PCR (20). *E. coli* expression clones were transformed into BL21(DE3) for expression. Plasmids for mammalian expression were made by subcloning Entry clones into transient transfection Destination vectors (pCAN-FLAG-DEST, pCAN-HA-DEST, and pEF-DEST51) using Gateway LR recombination per the manufacturer’s instructions.

### Protein production

*E. coli* proteins were expressed as described in (21). For insect cell expression, baculoviruses were generated by transfection of bacmid DNA into Sf9 cells as previously described (22). Titered viruses were used at an MOI of 3 to infect Tni-FNL cells (23) which were grown at 21 °C for 72 hours prior to harvest. Frozen insect cell pellets were thawed and homogenized at room temperature in 100 ml lysis buffer (Table S2) per liter of culture volume. Homogenized cell pellets were then lysed by passing through a Microfluidizer M-110EH (Microfluidics Corp.) at 7,000 psi. Lysate was clarified by ultracentrifugation at 104,600 × *g* for 40 min at 4 °C. Clarified lysate was filtered through 0.4 μM Whatman PES syringe filters (GE Lifesciences). All proteins were purified on NGC chromatography systems (BioRad Laboratories). Most proteins were purified as described for proteins with TEV-cleavable His6-MBP amino-terminal tags in (24) with modifications to buffers as noted in Table S2. Lysis buffers were the base buffers used for the initial IMAC step. Full-length Neurofibromin was produced with a construct containing a His6 tag which was not removed during the purification. This protein was purified by the same method as in (24) but with only a single IMAC purification step followed by preparative size exclusion chromatography on Superose-6 Increase (GE Healthcare) in a 16/60 column. For all final proteins, concentrations were determined by measuring sample absorbance at 280 nm (Nanodrop 2000C Spectrophotometer, Thermo Scientific) and using the extinction coefficient calculated from the amino acid sequence.

### Size Exclusion Chromatography and SEC-MALS

For analytical SEC, freshly purified or thawed protein was centrifuged at 9,300 × *g* for 10 min at room temperature in a benchtop centrifuge. Superose-6 Increase (GE Healthcare) or Superdex S-200 (GE Healthcare) in 10/300 GL columns were used on NGC chromatography systems. Columns were equilibrated with final buffers (Table S2) and 0.5 ml of protein sample was run at a flow-rate of 0.5 ml/min. For SEC-MALS, proteins were centrifuged at 9,300 × *g* for 10 min at room temperature. Superose-6 increase or Superdex S-200 in 10/300 GL columns were used on an Agilent 1200 LC system in-line with Wyatt Heleos (light scattering) and Wyatt Optilab T-Rex (refractive index) detectors to assess molecular weight of the proteins. Bovine serum albumin (Thermo-Fisher) was used as a standard. Astra software (Wyatt Technology, version 7.1.3) was used to collect and analyze the SEC-MALS data. Detailed information on buffers, injection rates, and flow rates can be found in Table S3.

### Sedimentation velocity analytical ultracentrifugation

Sedimentation velocity experiments were carried out at 30,000 rpm and 20°C on a Beckman Coulter ProteomeLab XL-I analytical ultracentrifuge and An50-Ti rotor following standard protocols (25). Samples of full-length Neurofibromin at various concentrations (0.05 to 2.2 μM) in 20 mM HEPES, pH 7.4, 300 mM NaCl, and 1 mM TCEP were loaded in 12-mm two-channel epon centerpiece cells. Sedimentation velocity data were collected using the absorbance (280 or 230 nm depending on the concentration) and interference optical detection systems. Time corrected (26) data were analyzed in SEDFIT 16.1 (27) in terms of a continuous c(s) distribution of sedimenting species using an s range of 0 to 30 with a linear resolution of 300 and a maximum entropy regularization confidence interval of 0.68. The solution density and viscosity were measured experimentally at 20°C on an Anton Paar DMA5000 density meter and Anton Paar AMVn viscometer, respectively. Protein partial specific volumes were calculated based on their composition in SEDNTERP (28). Samples of ABCD and CDEF domains were similarly studied at various loading concentrations at 40,000 rpm and 20°C. ABCD was studied in 500 mM NaCl, 20 mM HEPES.NaOH, pH 7.3, and 5 mM TCEP, whereas CDEF was studied in 300 mM NaCl, 20 mM HEPES.NaOH, pH 7.3, and 5 mM TCEP. Similarly, samples of ABC and DEF were studied at 50,000 rpm and 20°C in a solution buffer of 300 mM NaCl, 20 mM Tris.HCl, pH 8.5, and 5 mM TCEP. In all cases, data were analyzed in SEDFIT 16.1 as described above using a sedimentation coefficient resolution of 0.1 S and regularization confidence interval of 0.68. Equimolar mixtures of ABCD and EF domains, as well as ABC and DEF domains, were studied in 300 mM NaCl, 20 mM Tris.HCl, pH 8.5, and 5 mM TCEP at 50,000 rpm and 20°C. Samples in the 1 to 5 μM concentration range were mixed from stock solutions of the individual domains just before the experiment, and data were analyzed as described above.

### Sedimentation equilibrium analytical ultracentrifugation

Sedimentation equilibrium experiments were carried out to confirm the oligomeric state of full-length Neurofibromin. Samples in 20 mM HEPES, pH 7.4, 300 mM NaCl, and 1 mM TCEP were loaded at concentrations of 0.55, 1.1, and 2.2 μM in 12 mm six-channel epon centerpiece cells (130 μl). Sedimentation equilibrium experiments were conducted at 20°C and 3,000, 5,000, and 7,000 rpm on a Beckman Optima XL-A analytical ultracentrifuge and An50-Ti rotor following standard protocols (25), with absorbance data collected at 280 nm. Data were analyzed globally in terms of a single non-interacting species in SEDPHAT 14.0 to obtain the molar mass.

### Negative stain transmission electron microscopy

Freshly purified (full-length) or thawed (domains) proteins were centrifuged at 9,300 × *g* for 10 min at 4 °C and diluted to 0.01 mg/ml in 20 mM Tris-Cl, pH 8.0, 300 mM NaCl, 5 mM TCEP. Diluted protein was adsorbed on freshly glow-discharged carbon-film grid (CF200-CU, Electron Microscopy Sciences), washed with dilution buffer and stained with 0.75% w/v uranyl formate, pH 4.5, for 30 sec. Images of full-length Neurofibromin were collected at 67,000X magnification with EPU software from Thermo Fisher on a Tecnai T12 electron microscope equipped with a Gatan Ultra Scan camera, with images recorded at a pixel size of 1.77Å. Images of CDEF were collected at 100,000X magnification with SerialEM on a Tecnai T20 electron microscope equipped with an Eagle CCD camera, with images recorded at a pixel size of 2.19Å. Images of ABC, ABCD, and DEF domains were collected at 40,000X magnification on a Hitachi 7650 electron microscope operating at 80kV with an AMT digital camera (Advanced Microscopy Techniques).

### 2D classification and 3D reconstruction of Neurofibromin proteins

Negatively stained Neurofibromin particles were autopicked using EMAN2 e2boxer using a box size of 326 pixels (29). Reference free 2D classification was carried out using the ISAC procedure implemented in Sphire package (30). 41,951 particles were initially selected. Particles representing disordered NF1 2D class averages were separated and the remaining particles were reclassified using ISAC. The final 294 class averages (representative images shown in Fig. 2B) representing 30,588 particles were used for *ab initio* model reconstruction. This model was used as a reference for 3D refinement using meridian with C_2_ symmetry in Sphire. The final resolution of the reconstructed model was ~ 20 Å, as estimated by Fourier shell correlation (criterion of 0.5). All further analysis and visualization was performed using Chimera (31). The 3D variability analysis was done on the final refined model in Sphire. The final model data was submitted to wwPDB under the accession code EMD-20667. Approximately, 28,800 particles of the NF1-CDEF domain were auto-picked using EMAN e2boxer and 2D classification was performed with ISAC procedure using the same approach as described above.

### Small-angle x-ray and neutron scattering data collection and analysis

Small-angle X-ray scattering (SAXS) experiments were performed on a Rigaku BioSAXS-2000 equipped with a Pilatus 100K detector (Rigaku Americas, The Woodlands, TX). A fixed sample-to-detector distance, calibrated with a silver behenate standard, was used to collect scattering profiles of Neurofibromin (0.5 and 1.0 mg/ml) at 20°C. The data was reduced with the instrument software to obtain the scattering intensity, *I*(*Q*), vs. wave vector transfer, *Q* (= 4π sin(θ)/λ, where 2θ is the scattering angle), and then buffer background subtracted.

Small-angle neutron scattering (SANS) experiments were performed on the Bio-SANS (CG-3) beamline at the High Flux Isotope Reactor (HFIR) located at Oak Ridge National Laboratory (ORNL). Measurements were made using a 15.5 m sample-to-detector distance for the main 1×1 m detector and 6 Å wavelength neutrons (32). The wing detector bank was utilized to capture scattering at high angles. Neurofibromin samples in H_2_O buffer (1 mg/ml) were loaded into 1 mm pathlength circular-shaped quartz cuvettes (Hellma USA, Plainville, NY) and measurements performed at 20°C. Data reduction followed standard procedures using MantidPlot (33). The measured scattering intensity was corrected for the detector sensitivity and scattering contribution from the solvent and empty cells, and then placed on an absolute scale using a calibrated standard (34).

The reduced SAXS and SANS data were analyzed using the ATSAS tools (35) PRIMUS and GNOM for Guinier and P(r) analysis, respectively. Reported zero-angle scattering intensity, I(0), and radius of gyration, R_g_, values were obtained from the P(r) analysis. The Neurofibromin molecular mass was calculated from I(0) and R_g_ of the SAXS data using the method of (36). The Neurofibromin mass was determined from I(0) of the SANS data as: *M* = *I*(0) *N*_*A*_/[*c*(Δ*ρ*)^2^*υ̅*^2^], where *N*_A_ = Avogadro’s number, *c* = protein concentration (= 1 mg/mL), Δ*ρ* = scattering length density contrast between protein and buffer solution (= *ρ*_prot_ – *ρ*_buf_), and *υ̅* = protein partial specific volume (= 0.74 mL/g), and. Using the Contrast module of MULCh (37), the NF1 sequence was used to calculate Δ*ρ* in H_2_O buffer (= 2.364 × 10^10^ cm^−2^). The DAMMIF program within ATSAS was used to generate *ab initio* shape reconstruction models to fit the Neurofibromin (0.5 mg/mL) SAXS data. A total of 20 models were generated and compared by the program. A theoretical SAXS curve was generated from the final negative-stained EM model of Neurofibromin holoprotein using the ATSAS tool EM2DAM.

### Immunoprecipitation and western blotting

An NF1-null HEK293T cell line was generated using CRISPR to knockout of both alleles of *NF1*. HEK293T and NF1-null 293T cells were maintained in DMEM with 10% fetal bovine serum. Transfection of plasmids into cells were performed using FuGENE 6 (Promega) according to the manufacturer’s protocol and the cells were harvested 24 hours after transfection. Cells were lysed in a lysis buffer containing 20 mM Tris-Cl, pH 7.5, 150 mM NaCl, 5 mM MgCl_2_, 1% Triton-X100, and protease inhibitor cocktail (Sigma).

Immunoprecipitation was performed using EZview Red Anti-FLAG M2 Affinity Gel (Sigma). Beads were incubated with the cell lysates for 2 hours at 4 °C and washed 3 times with cold lysis buffer. Primary antibodies (Cell Signaling Technology) used for western blotting were: FLAG (rabbit, 14793), V5 (rabbit, 13202), and HA (rabbit, 3724). LI-COR secondary antibodies were used and images were obtained using a LI-COR Odyssey scanner.

## RESULTS

### Neurofibromin forms a high-affinity dimer in vitro

Purified full-length human Neurofibromin produced in baculovirus-infected insect cells migrated on SDS-PAGE at a molar mass close to its predicted size of 317 kDa (Fig. 1A). However, the purified protein eluted from a size exclusion column at a position equivalent to the 669 kDa thyroglobulin standard (Fig. 1B) suggesting that purified Neurofibromin existed as a dimer. To confirm this, additional measurements were made including SEC-MALS (size exclusion chromatography - multi-angle light scattering) which calculated a molar mass of 644 kDa (Fig. 1C), again consistent with a dimeric form of the protein. Additionally, we used small-angle X-ray scattering (SAXS) and small-angle neutron scattering (SANS) to investigate the structure and oligomeric state of Neurofibromin in solution (Fig. 1D). SAXS was performed at two Neurofibromin concentrations (0.5 and 1.0 mg/ml) with both yielding similar results, and SANS was also used to obtain lower Q data (< 0.01 Å^−1^). The protein molar mass predicted from SAXS and SANS were in agreement, consistent with a dimeric architecture for Neurofibromin (Table S1). Finally, analytical ultracentrifugation was used to conclusively characterize the oligomeric state of full-length Neurofibromin across a range of concentrations. Analysis of sedimentation velocity and sedimentation equilibrium experiments at various concentrations ranging from 1.2 to 0.05 μM showed the presence of a single dominant species across the entire concentration range which sedimented at 15.04 S with an estimated molar mass of 610 kDa (Fig. 1E) and a best-fit molar mass of 620 ± 70 kDa (Fig. S1) respectively. Overall, these data strongly supported the conclusion that full-length Neurofibromin forms a high-affinity (K_dim_ < 5 nM) dimer *in vitro*. To confirm this interaction *in vivo*, differentially epitope tagged NF1 constructs (FLAG and V5 epitopes) were cotransfected into HEK293T cells. Antibodies against the FLAG epitope were used to immunoprecipitate (IP) FLAG-tagged Neurofibromin and precipitated samples were probed with anti-FLAG and anti-V5 antibodies (Fig. 1F). V5-tagged Neurofibromin was readily detected in the FLAG IP sample, providing evidence of an interaction between full-length Neurofibromin proteins in cells, consistent with dimerization.

**Figure 1.**
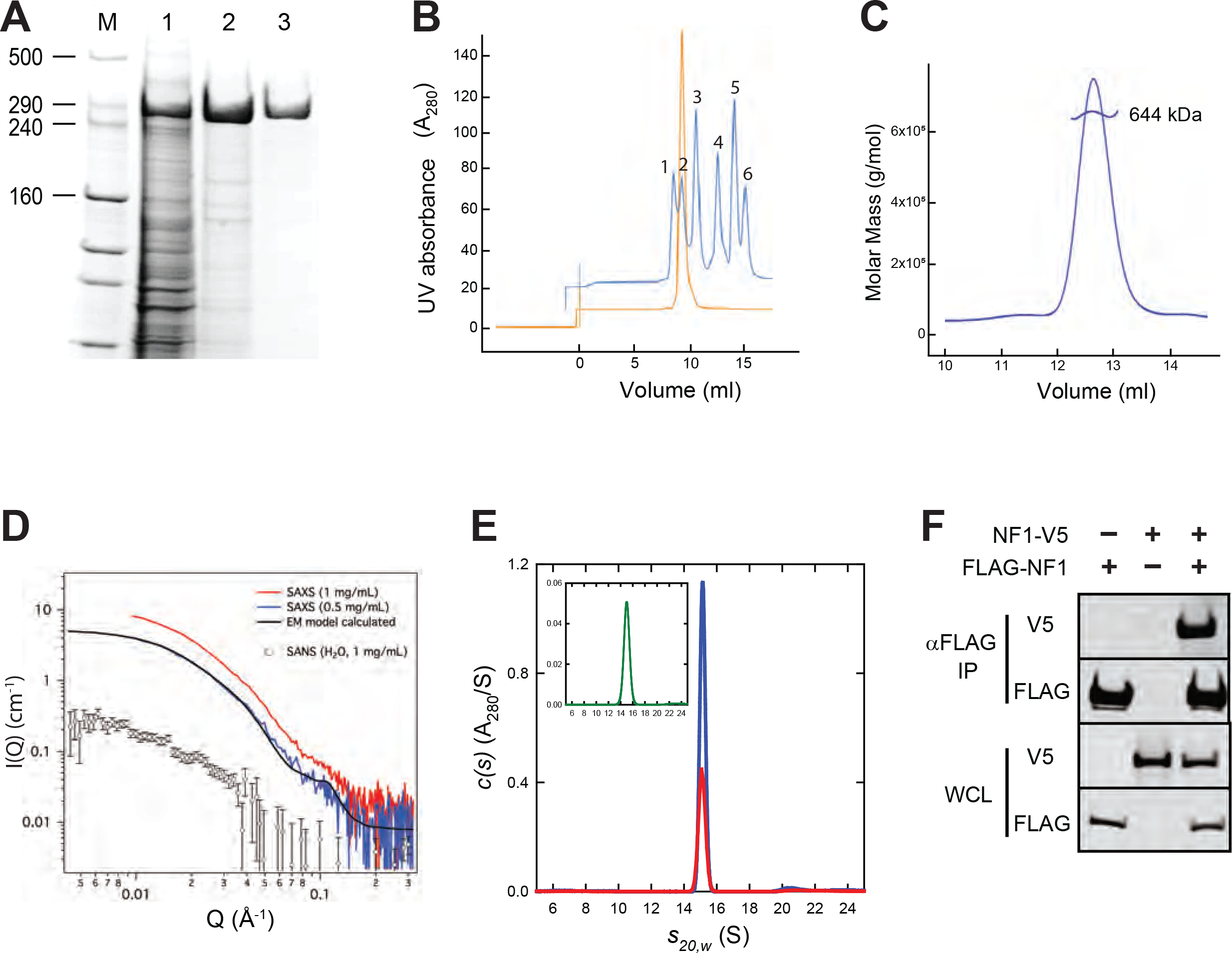
Full-length Neurofibromin is a high-affinity dimer. *A*, SDS-PAGE analysis of purified full-length Neurofibromin. Lanes are molecular weight markers (M) with sizes of relevant bands shown in kDa, clarified extract from the insect cell expression (*1*), elution pool from IMAC chromatography (*2*), and purified protein after size-exclusion chromatography (*3*). *B*, analytical SEC trace of purified Neurofibromin (orange) compared with molecular weight standards (blue). Standards used were blue dextran (*1*), thyroglobulin (*2*, 669 kDa), ferritin (*3*, 440 kDa), aldolase (*4*, 158 kDa), conalbumin (*5*, 75 kDa), and ovalbumin (*6*, 43 kDa). *C*, SEC-MALS analysis of full-length Neurofibromin. *D*, SAXS/SANS analysis of full-length Neurofibromin. Red (1 mg/ml) and blue (0.5 mg/ml) lines are SAXS data from runs with two different concentrations of Neurofibromin, while open circles represent SANS data using 1 mg/ml Neurofibromin. The black line represents theoretical SAXS data predicted by the 3D EM model. *E*, sedimentation velocity absorbance c(s) profiles for NF1 at 0.6 μM (red) and 1.2 μM (blue) based on data collected at 280 nm. The inset shows the corresponding c(s) profile for NF1 at 50 nM (green) based on absorbance data collected at 230 nm. *F*, Western blot of co-immunoprecipitation of differentially epitope-tagged NF1 proteins in HEK293 cells. The top two gel sections contain lysate purified with anti-FLAG antibodies. The bottom two sections contain whole-cell lysate (WCL). In both cases, the samples are probed with antibodies to the FLAG or V5 epitopes as noted.

**Figure 2.**
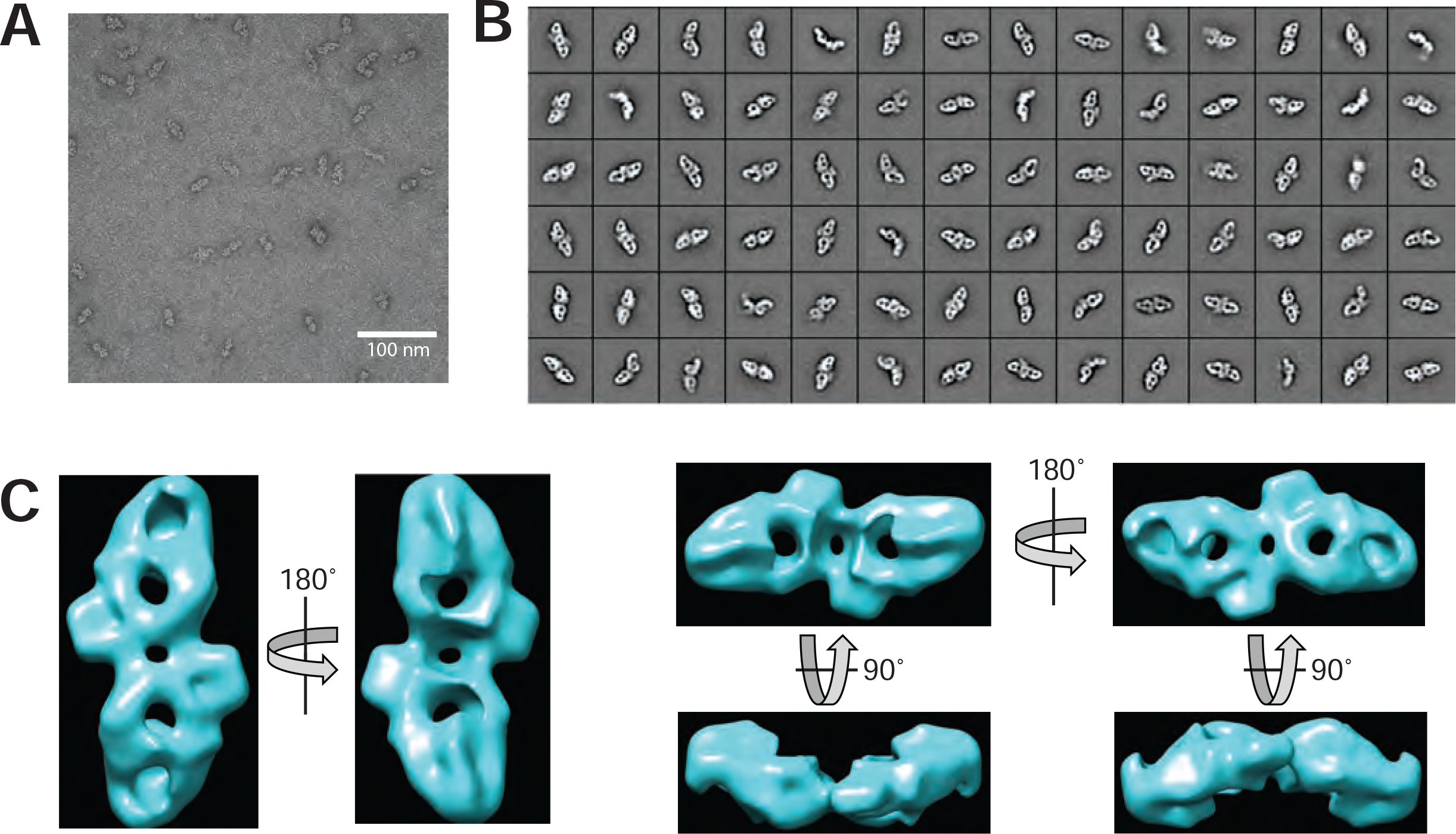
Negative-stain electron microscopy of full-length Neurofibromin. *A*, representative transmission electron micrograph of full-length Neurofibromin protein. *B*, representative 2D class averages obtained from 41,951 particles. *C*, 20 Å density map of full-length Neurofibromin generated from 294 class averages.

### Negative stain Electron Microscopy (EM) shows the dimeric structure of Neurofibromin

Projection images of negatively stained Neurofibromin obtained by transmission electron microscopy analysis allowed visualization of the Neurofibromin particle as a pseudo-symmetric dimer (Fig. 2A, 2B) in which the two NF1 protomers appear as oblong lobes connected via an interface region. Inspection of the 2D class averages suggest the dimeric arrangement is conformationally heterogeneous, with indications of variability both in quaternary structure and in the individual protomers. 3D reconstruction of the structure at ~ 20 Å resolution (Fig. 2C) using the dominant 2D classes allows visualization of the densities corresponding to the two NF1 protomers. The complex has overall dimensions of ~32 nm along the long axis of the dimer. Back projections from this model using EMAN2 to generate 2D class averages were used to validate the accuracy of the model (Fig. S2). 3D variability analysis (Fig. S3) highlights the unreliable parts of the reconstruction corresponding to regions of diffuse density seen in some 2D class averages. These data support the notion of some level of conformational flexibility in portions of the Neurofibromin dimer. A theoretical scattering curve generated from the reconstructed 3D model (Fig. 1D, black line) shows good agreement with the experimental SAXS and SANS data. Additionally, the overall size and shape of the structure seen in the 3D reconstruction is consistent with the solution structure of the protein predicted from *ab initio* SAXS/SANS data (Fig. S4). The SANS prediction of a maximum dimension (D_max_) in solution of ~30 nm is in reasonable agreement with predictions from the negative stain data and suggests that the reconstruction may be slighter larger than the actual solution size due to flattening or dehydration effects during EM grid preparation.

### Amino-terminal domains of Neurofibromin are incapable of dimerization

To investigate the elements of Neurofibromin that might be required for dimerization, we divided the protein into a series of domains (A through F, Fig. 3) based on bioinformatic analysis including phylogenetic homology, intron/exon boundaries, secondary structure prediction, and surface entropy prediction. DNA constructs representing the various combinations of domains were generated that corresponded to 21 different protein fragments. A subset of these proteins were purified and used for the experiments discussed in this manuscript (Table 1). Amino-terminal ABC and ABCD fragments could be purified at high yield and were utilized for biochemical analysis (Fig. 4A). Under reducing conditions, both ABC and ABCD migrated on SDS-PAGE near their expected molar masses (173 kDa and 206 kDa, respectively). SEC-MALS produced sharp peaks corresponding to molar masses of 174 and 211 kDa, respectively (Fig. 4D), suggesting that these proteins exist in solution as monomers. This result was confirmed using analytical ultracentrifugation, in which sedimentation velocity of the ABC domain at multiple concentrations (Fig. 4B) showed the presence of a species sedimenting at 6.88S with an estimated molar mass of 165 kDa, consistent with the size of an ABC monomer. Similar observations were made with ABCD at 1.1 μM and showed the presence of a species at 7.58 S with an estimated molar mass of 190 kDa (Fig. 4C), confirming that the ABCD domain is also monomeric. The smaller than expected mass can be attributed to the weak self-association observed as the concentration was raised from 1.1 to 4.4 μM. SAXS data also confirmed that the ABCD domain was monomeric in solution (Table S1, Fig. S5). Transmission EM studies of the ABC and ABCD proteins showed elongated particles of a monomeric size which could not be consistently class-averaged, suggesting a flexible structure (Fig. S6, panels A and B). Similar domain boundaries were used to engineer constructs for expression in HEK293T cells in which endogenous NF1 was deleted on both alleles. Differentially epitope labeled ABC and ABCD proteins were unable to co-immunoprecipitate (co-IP), confirming the lack of interaction of these monomeric domains *in vivo* (Fig. 4E, green circled region).

**Figure 3.**
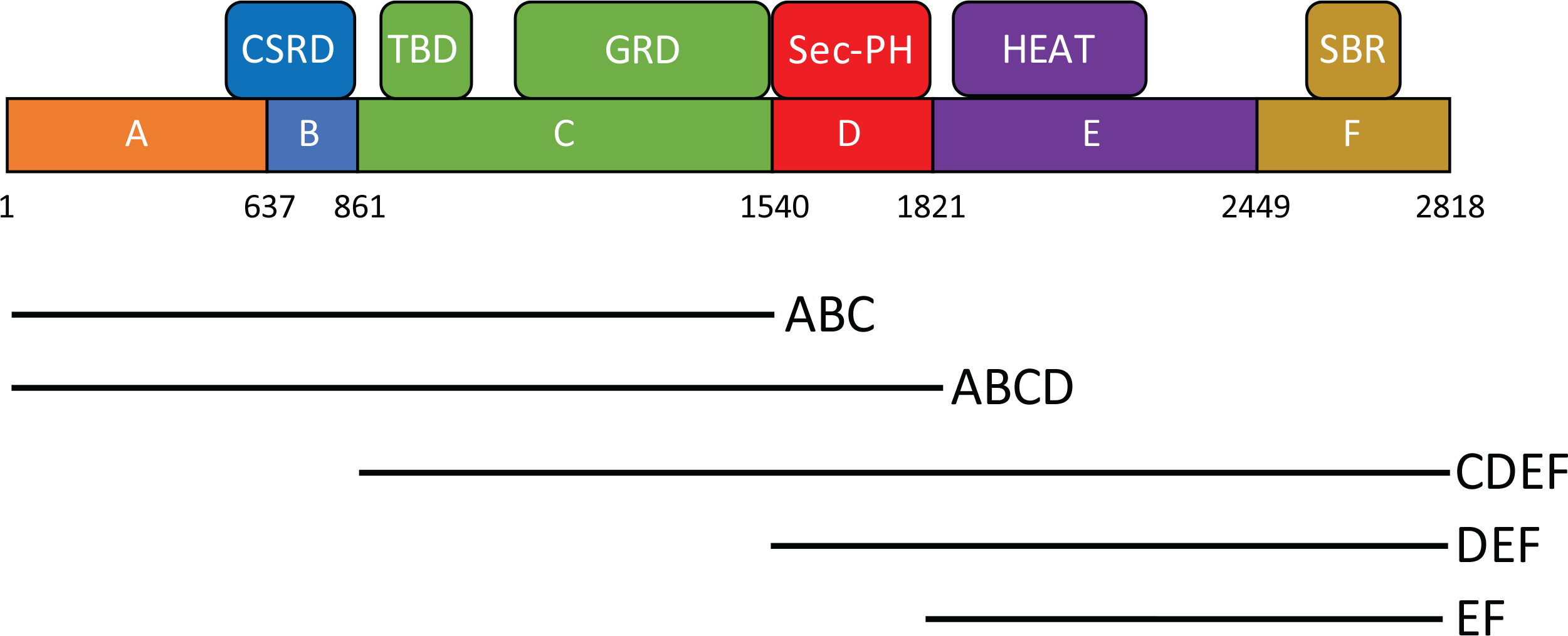
Neurofibromin fragments and domain architecture. Schematic of the regions utilized for production of Neurofibromin proteins in this manuscript. Domains have amino acid boundaries as noted and are designated by letters A through F. Corresponding regions of interest identified from the literature are noted above. Abbreviations are CSRD, cysteine-serine rich domain; TBD, tubulin-binding domain; GRD, GAP-related domain; Sec-PH, Sec14-pleckstrin homology; HEAT, HEAT-like repeat domain; SBR, syndecan-binding region.

**Figure 4.**
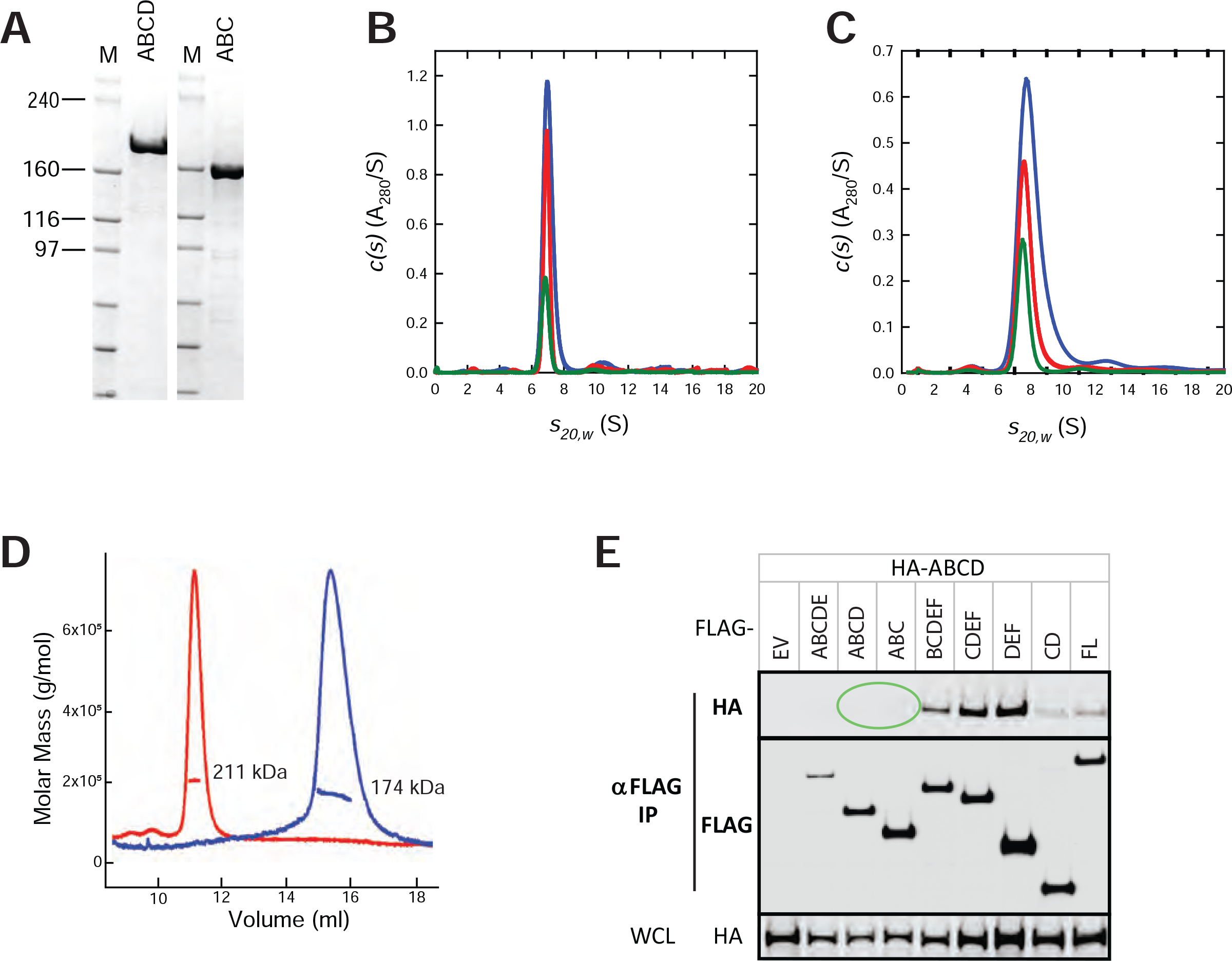
Amino-terminal domains of Neurofibromin are monomeric. *A*, SDS-PAGE gel of purified ABC and ABCD proteins. Size of molecular weight markers are noted in kilodaltons. *B*, sedimentation velocity absorbance c(s) profiles for the ABC protein at 1.1 μM (green), 2.2 μM (red), and 4.5 μM (blue) based on data collected at 280 nm. *C*, sedimentation velocity absorbance c(s) profiles for the ABCD protein at 1.1 μM (green), 2.1 μM (red), and 4.2 μM (blue) based on data collected at 280 nm. *D*, SEC-MALS analysis of ABC (blue) and ABCD (red) fragments. *E*, Western blot of co-immunoprecipitation of differentially epitope-tagged NF1 proteins in HEK293 cells. In this figure, HA-ABCD protein was co-IPed with FLAG-tagged domains as noted. The top two gel sections are lysates purified with anti-FLAG antibodies and probed with antibodies to the HA or FLAG epitopes. The bottom section contains whole-cell lysate (WCL) probed with anti-HA antibodies. Circled regions are discussed in more detail in the text. EV, empty vector control; FL, full-length NF1.

**Table 1.** Neurofibromin proteins used in this manuscript. Cited below are the amino acid regions present in purified protein domains, along with predicted isoelectric point (pI) and predicted molecular weight (MW) of each final protein. Also shown is the construct from which the proteins were purified and the host organism used for protein expression. In the case of His6-MBP fusion proteins, the final purified proteins were cleaved from the solubility tag, while full-length His6-tagged Neurofibromin remained tagged after purification. Constructs C* and C*D have altered start sites within the C domain compared with other C domain containing proteins.

| Domain | Amino Acids | pI | MW (kDa) | Exp. Host | Construct |
| --- | --- | --- | --- | --- | --- |
| Full length | 2-2818 | 6.9 | 317 | Tni-FNL | His6-tev-NF1 |
| ABC | 2-1540 | 6.4 | 173 | Tni-FNL | His6-MBP-tev-NF1(2-1540) |
| ABCD | 2-1820 | 6.5 | 206 | Tni-FNL | His6-MBP-tev-NF1(2-1820) |
| CDEF | 861-2818 | 6.9 | 220 | Tni-FNL | His6-MBP-tev-NF1(861-2818) |
| DEF | 1541-2818 | 8.0 | 144 | Tni-FNL | His6-MBP-tev-NF1(1541-2818) |
| EF | 1821-2818 | 8.1 | 111 | Tni-FNL | His6-MBP-tev-NF1(1821-2818) |
| C*D | 1085-1816 | 6.4 | 83 | Tni-FNL | His6-MBP-tev-NF1(1085-1816) |
| D | 1541-1820 | 7.3 | 32 | Tni-FNL | His6-MBP-tev-NF1(1541-1820) |
| C* | 1085-1530 | 6.0 | 50 | Tni-FNL | His6-MBP-tev-NF1(1085-1530) |
| GRD | 1172-1530 | 6.0 | 41 | *E. coli* | His6-MBP-tev-NF1(1172-1530) |
| GRD | 1198-1530 | 6.4 | 38 | *E. coli* | His6-MBP-tev-NF1(1198-1530) |
| GRD | 1203-1530 | 6.6 | 38 | *E. coli* | His6-MBP-tev-NF1(1203-1530) |

### Carboxy-terminal domains of Neurofibromin are capable of dimerization

As the amino-terminal end of Neurofibromin did not appear capable of forming dimers, we next examined the carboxy-terminal end of the protein using fragments CDEF, DEF, and EF. All three polypeptides could be produced and purified at high yield (Fig. 5A), and migrated near their expected molar masses on SDS-PAGE (220 kDa, 144 kDa, and 111 kDa, respectively). SEC-MALS data, however, showed anomalous migration of these proteins, yielding molar masses of 345 kDa (CDEF), 273 kDa (DEF), and 204 kDa (EF), suggesting that these proteins were capable of self-association into higher-order structures (Fig. 5D). Sedimentation data for DEF collected at 1.1 μM showed the presence of a species at 8.52 S with an estimated mass of 170 kDa. When the concentration was increased to 10.5 μM, a species was observed at 9.56 S (Fig. 5B). The larger than monomer mass observed for the 8.52 S species at 1.1 μM, the observation of intermediate sedimentation coefficient species at concentrations between 1.1 and 10.5 μM, and the observation of a best-fit frictional coefficient of less than unity at 10.5 μM support a weak self-association with fast-exchange (38). Sedimentation experiments on CDEF at 8.8 μM showed the presence of a species at 11.65 S with an estimated molar mass of 300 kDa (Fig. 5C). This data suggests that the CDEF domain is dimeric and the smaller than expected mass can be attributed to the self-association observed as the concentration is raised from 2.2 to 8.8 μM. SAXS data on CDEF at 15 uM also confirmed that this protein was entirely dimeric in solution at high concentration (Table S1, Fig. S5). The oligomerization of this domain is strongly concentration dependent; based on SAXS and AUC data, CDEF exists as a dimer at concentrations above 150 nM. At the lower concentrations used for negative stain EM analysis (45 nM), a mixed population of monomers and dimers were observed (Fig. S6C).

**Figure 5.**
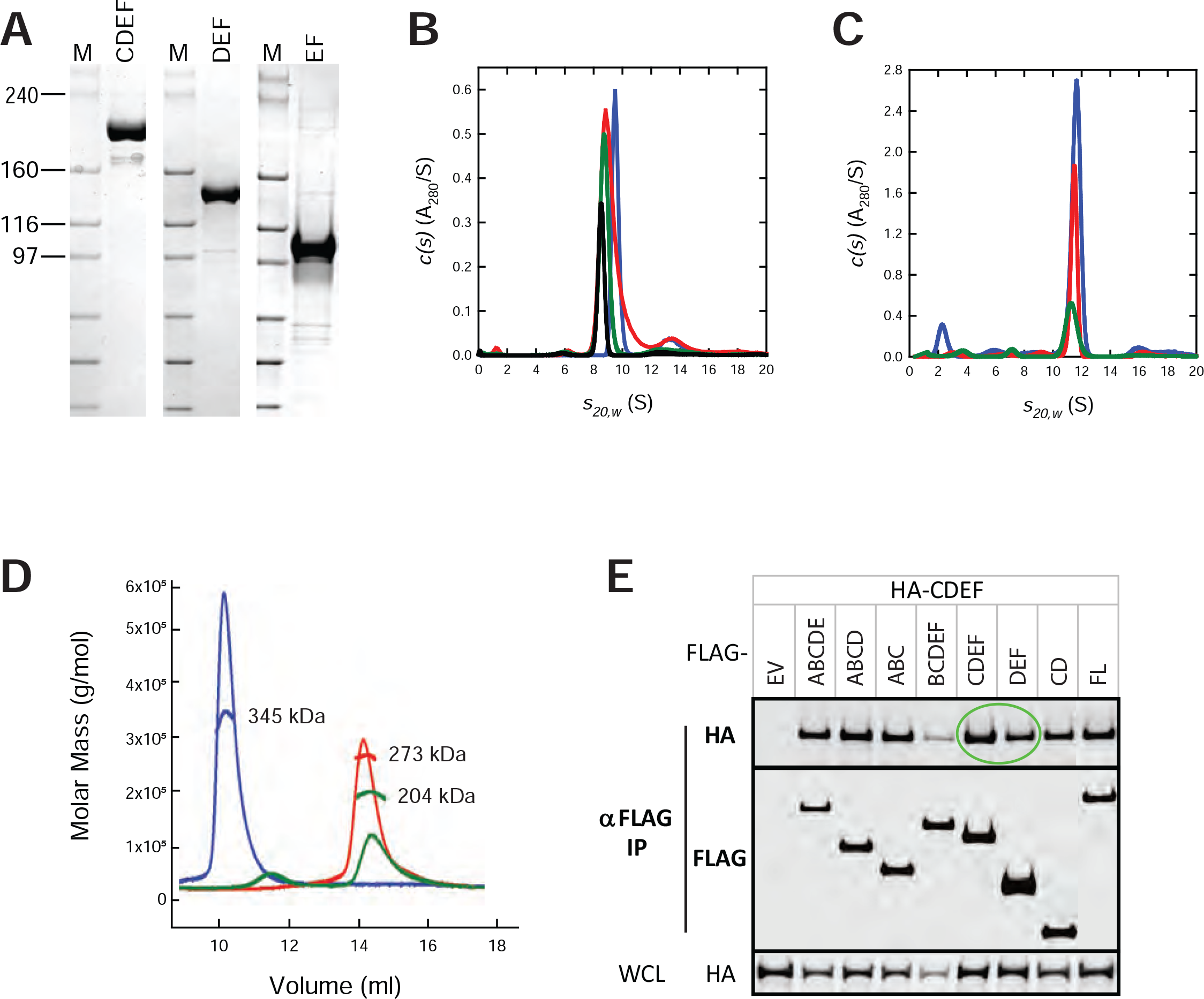
Carboxy-terminal domains of Neurofibromin are capable of dimerization. *A*, SDS-PAGE gel of purified CDEF, DEF, and EF proteins. Size of molecular weight markers are noted in kilodaltons. *B*, sedimentation velocity absorbance c(s) profiles for the CDEF protein at 2.1 μM (green), 4.2 μM (red), and 8.8 μM (blue) based on data collected at 280 nm. *C*, sedimentation velocity absorbance c(s) profiles for the DEF protein at 1.2 μM (black), 2.6 μM (green), 5.2 μM (red), and 10.5 μM (blue) based on data collected at 280 nm. Data at the highest concentration was collected using a 3 mm pathlength cell, standard 12 mm cells were used for the lower concentrations. *D*, SEC-MALS analysis of CDEF (blue), DEF (red), and EF (green) proteins. *E*, Western blot of co-immunoprecipitation of differentially epitope-tagged NF1 proteins in HEK293 cells. In this figure, HA-CDEF protein was co-IPed with FLAG-tagged domains as noted. The top two gel sections are lysates purified with anti-FLAG antibodies and probed with antibodies to the HA or FLAG epitopes. The bottom section contains whole-cell lysate (WCL) probed with anti-HA antibodies. Circled regions are discussed in more detail in the text. EV, empty vector control; FL, full-length NF1.

Finally, using HEK293 cells lacking endogenous NF1, we were able to show that the CDEF and DEF domains of differentially epitope-tagged Neurofibromin could co-IP with each other (Fig. 5E, green circled region), confirming the potential for dimerization in cells similar to that observed with purified proteins.

### Equimolar mixtures of amino-terminal and carboxy-terminal domains of Neurofibromin can reconstitute the full-length Neurofibromin dimer

The ability of CDEF to self-associate but not form dimers at concentrations observed for the full-length protein, suggests that the subdomains may lack critical intramolecular or intermolecular contacts that stabilize the dimer. In order to test this hypothesis, purified ABC and DEF proteins were mixed in an equimolar concentration and subjected to size exclusion chromatography. Surprisingly, the major peak of protein was observed eluting from the SEC column at a size corresponding to 600 kDa, similar to that observed with full-length Neurofibromin dimers. SDS-PAGE analysis confirmed that both subdomains were present in this complex at what appeared to be stoichiometric levels (Fig. 6A). Sedimentation velocity experiments were carried out on the purified complex of ABC and DEF. Complexes made from an equimolar mixture of ABC and DEF in the 1.0 to 4.5 μM concentration range showed the presence of a species at 14.32 S with an estimated mass of 580 ± 70 kDa (Fig. 6D). We observed similar reconstitution results using ABCD and EF proteins and sedimentation studies of these mixtures in the 1.0 to 5.5 μM concentration range also showed the presence of a species at 14.26 S with an estimated mass of 580 ± 95 kDa (data not shown). Molar masses represent the average from all experiments, where the reconstituted dimer represents approximately 65% and 85% of the sedimenting signal for ABC/DEF, and ABCD/EF, respectively. Sedimentation data over a range of concentrations suggests that these complexes have similar dimerization affinities (<5 nM) as the full-length Neurofibromin dimers. Electron microscopic images and resulting 2D class averages show the presence of structures in the reconstituted complexes that are similar to those observed with full-length Neurofibromin dimers (Fig. 6B and 6C). These results suggest that a mixture of monomeric N-terminal fragments such as ABC with C-terminal fragments capable of dimerization such as DEF can recapitulate full-length dimeric Neurofibromin structures with very high affinity. Co-IP data in HEK293 cells (Fig. 6E) also confirms that ABC and DEF proteins are capable of interactions in cells which permits pull-down of differentially tagged proteins—here, both amino-terminal fragments (ABC, ABCD) are capable of co-IP with the carboxy-terminal DEF fragment as predicted by the *in vitro* protein reconstitution data, as also shown in Fig. 5E for the CDEF fragment.

**Figure 6.**
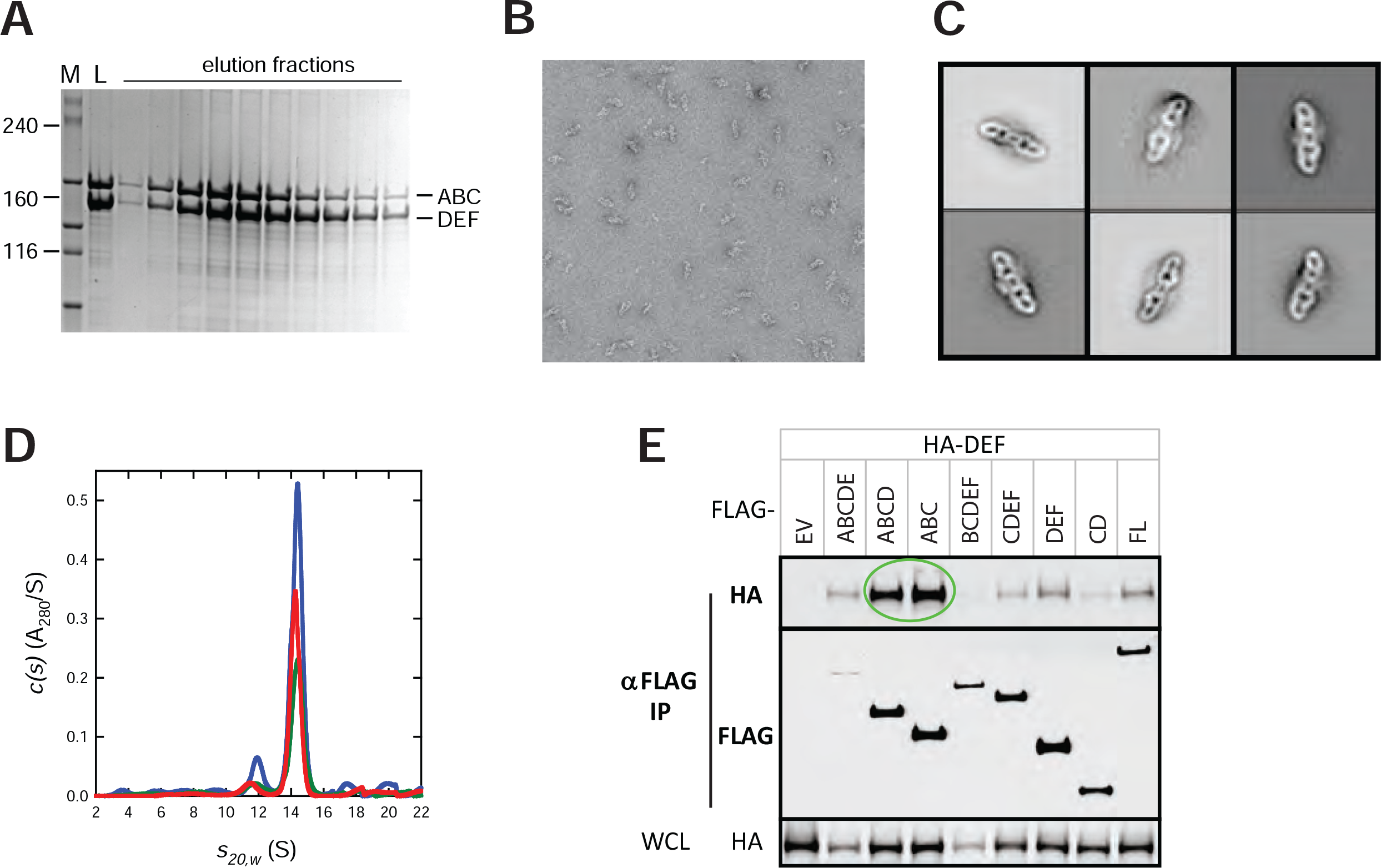
Reconstitution of full-length Neurofibromin dimers from ABC and DEF proteins. *A*, SDS-PAGE analysis of SEC of a equimolar mixture of ABC and DEF proteins. Size of molecular weight markers in lane M are noted in kilodaltons. Lane L represents the loaded material, and additional lanes are elution fractions across the column. *B*, transmission electron micrograph of reconstituted Neurofibromin dimers from the ABC/DEF mixture. *C*, representative 2D-class averages from particles selected from multiple TEM micrographs. *D*, sedimentation velocity absorbance c(s) profiles for equimolar mixtures of the amino-terminal ABC and carboxy-terminal DEF fragments of NF1 at 1.0 μM (green), 2.0 μM (red), and 4.5 μM (blue) based on data collected at 280 nm. Data at the highest concentration was collected using a 3 mm pathlength cell, standard 12 mm cells were used for the lower concentrations. *E*, Western blot of co-immunoprecipitation of differentially epitope-tagged NF1 proteins in HEK293 cells. In this figure, HA-DEF protein was co-IPed with FLAG-tagged domains as noted. The top two gel sections are lysates purified with anti-FLAG antibodies and probed with antibodies to the HA or FLAG epitopes. The bottom section contains whole-cell lysate (WCL) probed with anti-HA antibodies. Circled regions are discussed in more detail in the text. EV, empty vector control; FL, full-length NF1.

### Identification of regions responsible for Neurofibromin dimerization

In order to narrow down the regions responsible for dimerization, we investigated interactions between the C-terminal EF domain and regions upstream in Neurofibromin. We measured interactions using SEC by mixing the EF domain (1821-2818) with either the C*D domain (1085-1816, Fig. 7A) or the C* domain (1085-1530, Fig. 7B), which show co-elution indicating complex formation, while the D domain (1541-1820, Fig. 7C) does not. Using additional NF1 GAP-related domain (GRD) proteins, we further defined that the region between amino acids 1085-1172 was essential for interaction with the EF domain (Fig. S7). We were also able to confirm that the same region was essential for interaction with the DEF domain (Fig. S8), suggesting that this region, known as the tubulin-binding domain (TBD), is important for the multimerization of Neurofibromin.

**Figure 7.**
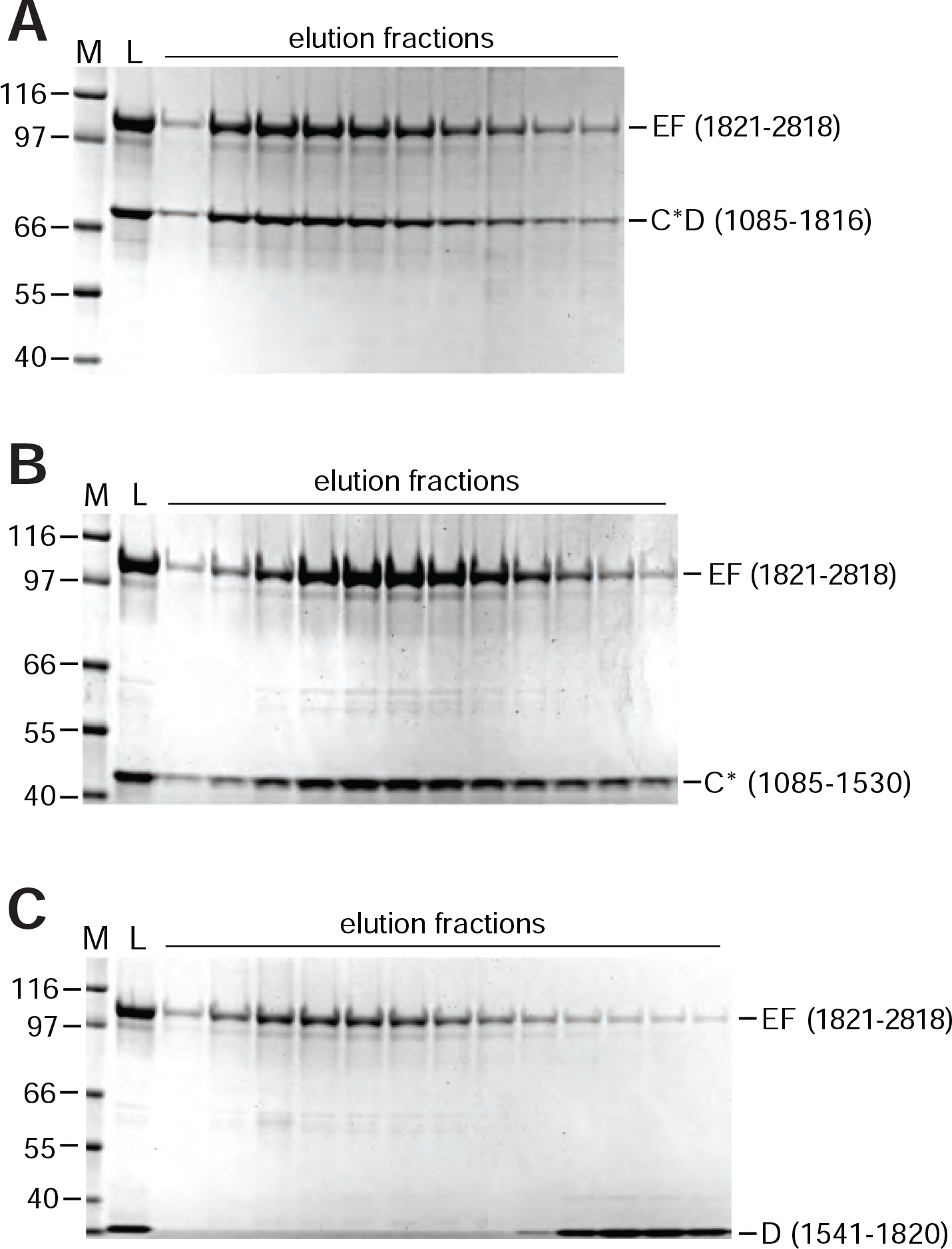
Identification of the interaction between the Neurofibromin C and EF domains. SDS-PAGE analysis of SEC of equimolar mixtures of NF1 domains with EF (1821-2818). In each panel, the size of molecular weight markers in lane M is noted in kilodaltons. Lane L represents the loaded material, and additional lanes are elution fractions across the column. Neurofibromin fragments mixed with the EF domain in each panel were: *A*, 1085-1816; *B*, 1085-1530; *C*, 1541-1820.

## DISCUSSION

The 2818 amino acid Neurofibromin protein is involved in a variety of human diseases including several sporadic cancers, as well as the common genetic disorder Neurofibromatosis type I. While the underlying mechanism of these diseases seems to involve defects in the GTPase-activating activity of Neurofibromin, which usually regulates the RAS family of small GTPases and downstream MAP kinase pathways, this activity resides in a small portion of the protein. Very little is understood about the role of any of the remaining parts of Neurofibromin. Also, missense mutations which lead to NF1 disease and NF1-driven cancers occur throughout the protein and are not localized specifically to the GAP-related domain (39). This argues for additional roles of these mutations in either Neurofibromin activity or protein stability. In order to investigate some of these questions, we produced full-length human Neurofibromin protein using an insect cell system. Size exclusion chromatography experiments showed that this large protein appeared to exist in solution as a dimer. Additional biophysical measurements confirmed that the purified Neurofibromin protein was dimeric and that the dimerization affinity was low enough that effectively all of the protein existed in solution as an obligate dimer at concentrations above 50 nM. We were able to confirm using cell-based methods that Neurofibromin proteins expressed in HEK293 cells lacking endogenous NF1 also showed self-association, as differentially tagged NF1 genes could be used to co-IP Neurofibromin dimers containing both tags. The dimeric organization of the protein was also verified in projection electron microscopic images and the resulting 3D reconstruction of purified Neurofibromin. These findings are consistent with a report showing that Ira2, one of the two *S. cerevisiae* paralogs of Neurofibromin, could coimmunoprecipitate the other paralog, Ira1, as detected by mass spectrometry (40). During the preparation of this manuscript, researchers noted that Neurofibromin eluted from their size exclusion chromatography during purification as an apparent dimer (41), and other studies have proposed that multimerization could potentially explain dominant-negative effects of heterozygous truncation mutations of NF1 in patients (42).

To define the domain architecture of Neurofibromin, we chose a directed bioinformatic approach in which the protein was broken down into six domains based on a series of protein-focused bioinformatic methods, and the various contiguous combinations of these domains were cloned, and proteins generated using a baculovirus-based insect cell system. Previous attempts to generate Neurofibromin fragments using a random library approach in *E. coli* generated few useful proteins (43). However, in our study, with few exceptions, we were able to purify most of the Neurofibromin fragment proteins using these domain boundaries. We focused on the well-purified amino-terminal domains (ABC, ABCD) and carboxyterminal domains (CDEF, DEF, EF) for additional studies.

Amino-terminal domains of Neurofibromin (ABC, ABCD) produced monomeric proteins at all concentrations studied. Negative stain EM images of these domains suggested the presence of partially unfolded particles. In contrast, the carboxy-terminal domains (CDEF, DEF, EF) produced proteins which were capable of self-association in a concentration dependent manner. In particular, the DEF and EF proteins showed very weak self-association which was evident only at the highest concentrations. However, the CDEF protein formed significant amounts of dimeric protein, and even at the low concentration used for EM analysis, a clear mixture of monomer and dimer populations was observed. This suggests that CDEF likely contains the major components responsible for the dimerization of the full-length protein. *In vivo* data using these domains confirmed these findings—ABC and ABCD proteins which were differentially tagged were unable to co-IP with each other *in vivo*, confirming that these proteins do not interact to form multimers. However, co-IP was observed with differentially tagged CDEF and DEF proteins, suggesting again that the experiments with purified proteins recapitulate what is observed in cells.

The *in vivo* data also produced an unexpected result —the ABC protein *in vivo* was able to co-IP with the DEF protein. In order to investigate whether this phenomenon could be recapitulated *in vitro*, we mixed the purified ABC and DEF proteins and separated the mixture using size exclusion chromatography. Surprisingly, a protein complex eluted from the SEC column at the size of full-length Neurofibromin dimer (600 kDa). This complex behaved in analytical ultracentrifugation experiments exactly like full-length dimers, including a lack of concentration dependence suggesting a high-affinity interaction as observed for the full-length protein. Projection electron microscopic images from negatively stained complexes were indistinguishable from those seen with the full-length protein, suggesting that this mixture of domains could recapitulate the folding of full-length Neurofibromin dimers. Similar reconstitution experiments were carried out with ABCD and EF proteins and produced identical results (data not shown).

Based on these results, we focused on potential interactions with the EF domain. The E domain of Neurofibromin contains a series of 12 predicted HEAT-like repeats. These repeats are commonly involved in protein-protein interactions, and often form a complex solenoid structure using a series of alpha-helices which can be used for additional interactions with partner proteins (44). From the reconstitution data, we theorized that an E domain solenoid might interact with a region upstream in Neurofibromin, likely in the C or D regions. To test this idea, we attempted to make complexes using purified C, D, or CD domains with EF. Both the C domain and CD domains were able to make complexes with EF, but the D domain was not, implicating the C domain (which contains the putative tubulin-binding domain, TBD, and the GAP-related domain, GRD) in the interaction. As several C domain constructs used for studies of the structure of the GAP domain of Neurofibromin were available to test, we were able to identify that only the region of the C domain from 1085-1171 was required for complex formation with EF. Similarly, we confirmed that this TBD region of Neurofibromin was able to form stable complexes with the DEF domain protein as well.

The Neurofibromin TBD consists of three predicted amphipathic alpha-helices (1095-1113, 1134-1152, 1173-1194) which we hypothesize might be capable of interacting with the E domain HEAT-like solenoid. This type of tight interaction involving a HEAT-like solenoid at a dimer interface has been observed with the adaptor protein mTOR. Cryo-EM studies have shown that the formation of high-affinity mTOR dimers is due to the hydrophobic interaction of a HEAT-like solenoid in the N-terminal region of mTOR with a set of three amphipathic helices in the core domain of a second monomer of mTOR (45). Like Neurofibromin, the mTOR dimer has an equally high affinity (10 nM) due to the strong interaction of these hydrophobic domains. The strength of these dimers is enhanced by the presence of multiple interactions, as each protomer contains both helical elements which leads to two large hydrophobic contact points along the dimer interface. We propose that the points of contact seen in the 3D reconstruction of Neurofibromin likely represent C-E domain interactions, leading to a stable, high-affinity dimer.

Dimerization provides a potential explanation for the phenotypes observed in common NF1 disease mutations. Many of the mutations observed are heterozygous nonsense or frameshift mutations, which lead to premature truncation of Neurofibromin. In most of these patients, there is still a single copy of wild type NF1 present, and yet as seen in a recent study, the total amount of full-length Neurofibromin in some of these patients is considerably less than predicted (*i.e.* 50% of wild type levels) (42). One explanation for this would be that the truncated forms of the protein are still capable of interactions (intermolecular or intramolecular) with the wildtype protein, which could lead to enhanced proteosomal degradation of the defective proteins. Effectively, these defective truncated proteins would serve as subunit poisons, still interacting with wildtype Neurofibromin and interfering with the production of normal dimers. The biophysical techniques utilized in this work should serve as an effective means of testing these mutants to see if their effects on wild type Neurofibromin can be recapitulated *in vitro*

The biological consequence of dimerization of Neurofibromin remains unclear. Interactions of RAS proteins with the NF1 GAP domain appear to involve monomers of both proteins, and the recently elucidated structure of KRAS-NF1 GAP-SPRED1 EVH also involves a 1:1:1 complex of monomeric proteins (Wupeng Yan, personal communication). It is possible that dimerization of Neurofibromin is required for other biological activities, or is in some way involved in the regulation of the protein. It also may be possible that Neurofibromin dimers are disrupted during the signaling process and that a transition to monomeric Neurofibromin is needed for interaction with RAS proteins. Additional work is needed to understand the possible biological ramifications of Neurofibromin dimerization.

## ACKNOWLEDGEMENTS

We thank members of the FNLCR Protein Expression Laboratory for help with cloning (Vanessa Wall, Jennifer Mehalko), E. coli expression (Troy Taylor, Nitya Ramakrishnan, Jose Sanchez Hernandez), insect cell expression (Kelly Snead), and purification (Peter Frank, Randy Gapud). We thank Giovanna Grandinetti, Joseph Darling, and Oleg Kuybeda for initial electron microscopic analysis. We also thank Andy Stephen, Dhirendra Simanshu, Lucy Young, and Claire Lorenzo for helpful discussions on the data in this manuscript. HON, CS, DB, and PJ acknowledge the support of the Laboratory Directed Research and Development Program of Oak Ridge National Laboratory (ORNL) for SAS and EM characterization. SANS studies on Bio-SANS were supported by the Office of Biological and Environmental Research (OBER) funded Center for Structural Molecular Biology (CSMB) under Contract FWP ERKP291, using the High Flux Isotope Reactor supported by the Office of Basic Energy Sciences (BES), U. S. Department of Energy (DOE). Sample preparation and SAXS studies were carried out using facilities at the Spallation Neutron Source at ORNL, supported by the DOE BES Scientific User Facilities. ORNL is managed by UT-Battelle, LLC, for the U. S. Department of Energy (DOE) under contract No. DE-AC05-00OR22725. This work has been funded in whole or in part with federal funds from the National Cancer Institute, National Institutes of Health, under contract HHSN261200800001E, and the Intramural Research Program of the National Institute of Diabetes and Digestive and Kidney Diseases (NIDDK). The content of this publication does not necessarily reflect the views or policies of the Department of Health and Human Services, nor does mention of trade names, commercial products, or organizations imply endorsement by the U.S. Government.

## Competing interests

The authors declare that they have no competing interests with the contents of this article.

## Author contributions

M.S., S-W. H., S.M., W.G., S.S., D.N., F.M., D.E. conceptualization; M.S., W.G., D.E. data curation; H.O., C.S., D.B., R.G. formal analysis; H.O., A.R., R.G., D.N., F.M., D.E. funding acquisition; M.S., S-W.H., M.D., P.J., D.B., C.S., R.G. investigation; M.S., S-W.H., S.M., M.D., W.G., P.J., H.O., C.S., D.B., R.G., S.S., D.E. methodology; M.S., S.M., W.G., P.J., H.O., A.R., R.G., D.E. project administration; M.S., S-W.H., S.M., M.D., P.J., C.S., D.B., R.G., S.S., F.M., D.E. resources; S.M., W.G., H.O., A.R., D.N., S.S., F.M., D.E. supervision; M.S., S-W.H., S.M., M.D., W.G., P.J., H.O., C.S., D.B., R.G., D.E. validation; M.S., S-W.H., W.G., P.J., H.O., C.S., D.B., R.G., D.E. visualization; W.G., H.O., C.S., R.G., D.E. writing-original draft.

---

**Table S1.** Neurofibromin protein size and mass predictions from SAXS and SANS analysis. Values are calculated from the data in Fig. 1D and Fig. S5, including radius of gyration (R_g_), maximum dimension (D_max_), predicted molecular mass (M) and predicted oligomeric state.

| Protein | Experiment | $I(0)^1$ | $R_g$ (Å) | $D_{max}$ (Å) | $M$ (kDa) | Oligomer ($M/M_{mon}$)<sup>2</sup> |
| --- | --- | --- | --- | --- | --- | --- |
| Full-length | SAXS (1.0 mg/ml) | $9.99 \pm 0.1$ | $89.4 \pm 0.9$ | 270 | $670 \pm 20$ | $2.11 \pm 0.06$ |
| Full-length | SAXS (0.5 mg/ml) | $4.73 \pm 0.08$ | $88.0 \pm 1.0$ | 258 | $630 \pm 20$ | $1.99 \pm 0.06$ |
| Full-length | SANS (1.0 mg/ml) | $0.302 \pm 0.004$ | $107.8 \pm 0.9$ | 300 | $593 \pm 8$ | $1.87 \pm 0.03$ |
| ABCD | SAXS (1.5 mg/ml) | $6.2 \pm 0.1$ | $67.1 \pm 0.7$ | 205 | $201 \pm 6$ | $0.98 \pm 0.03$ |
| CDEF | SAXS (4.3 mg/ml) | $6.6 \pm 0.1$ | $73.5 \pm 0.7$ | 228 | $450 \pm 10$ | $2.05 \pm 0.05$ |

<sup>1</sup>SAXS $I(0)$ values are in arbitrary units; SANS $I(0) = [\text{cm}^{-1}]$ .
<sup>2</sup> $M_{mon}$ , monomer molecular weight; 317 kDa (full length), 220 kDa (CDEF), 206 kDa (ABCD)

**Table S2.** Buffers used in protein purification. Cell pellets used to purify various Neurofibromin proteins were homogenized in the lysis buffers noted below prior to lysis by microfluidization. Lysis buffers also served as the base buffers for the IMAC purification steps. The final buffers were used as the running buffer during preparative size exclusion chromatography and for long-term storage of purified proteins. All lysis buffers also contained protease inhibitors (1:100 v/v, Sigma P-8849), and all final buffers contained protease inhibitors (1:1000 v/v, Sigma P-8849).

| Domain | Amino Acids | Lysis Buffer<sup>1</sup> | Final Buffer<sup>1</sup> |
| --- | --- | --- | --- |
| Full length | 2-2818 | A | A |
| ABC | 2-1540 | B | C |
| ABCD | 2-1820 | D | D |
| CDEF | 861-2818 | D | E |
| DEF | 1541-2818 | B | C |
| EF | 1821-2818 | C | C |
| C*D | 1085-1816 | C | C |
| D | 1541-1820 | F | G |
| C* | 1085-1530 | D | G |
| GRD | 1172-1530 | F | G |
| GRD | 1198-1530 | F | G |
| GRD | 1203-1530 | F | G |

<sup>1</sup>A: 20 mM Tris-Cl, pH 8.0, 300 mM NaCl, 5 mM TCEP
B: 20 mM Tris-Cl, pH 8.5, 500 mM NaCl, 10% (v/v) glycerol, 5 mM TCEP
C: 20 mM Tris-Cl, pH 8.5, 300 mM NaCl, 5 mM TCEP
D: 20 mM HEPES, pH 7.3, 500 mM NaCl, 5 mM TCEP
E: 20 mM HEPES, pH 7.3, 300 mM NaCl, 5 mM TCEP
F: 20 mM HEPES, pH 7.3, 300 mM NaCl, 1 mM TCEP
G: 20 mM HEPES, pH 7.3, 150 mM NaCl, 1 mM TCEP

**Table S3.** SEC-MALS Buffers/Conditions. All running buffers contained 20 mM Tris-Cl at the indicated pH along with 300 mM NaCl and 5 mM TCEP.

| Domain | SEC Resin | Running Buffer pH | Injection Volume (μl) | Protein Load (μg) | Flow Rate (ml/min) |
| --- | --- | --- | --- | --- | --- |
| Full length | Superose-6 Increase | 8.0 | 100 | 100 | 0.5 |
| ABC | Superose-6 Increase | 8.5 | 100 | 100 | 0.3 |
| ABCD | Superdex S200 | 8.0 | 50 | 127 | 0.3 |
| CDEF | Superdex S200 | 8.0 | 50 | 100 | 0.3 |
| DEF | Superose-6 Increase | 8.5 | 50 | 65 | 0.3 |
| EF | Superose-6 Increase | 8.5 | 100 | 80 | 0.3 |

**Figure S1.**
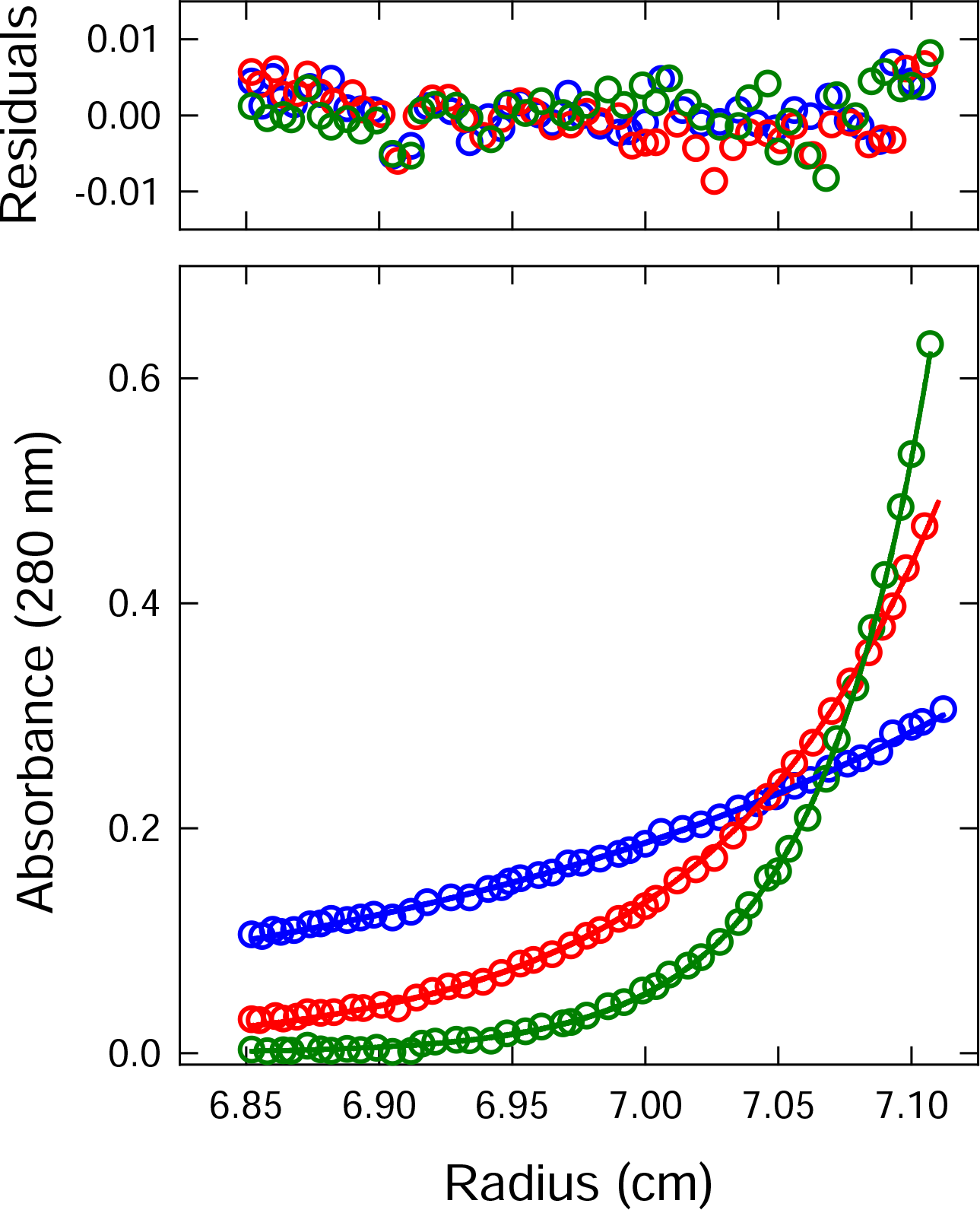
Sedimentation equilibrium of full-length Neurofibromin. Sedimentation equilibrium absorbance data for a solution of Neurofibromin at 550 nM collected at 3,000 (blue), 5,000 (red) and 7,000 (green) rpm analyzed in terms of a non-interacting single ideal solute. Solid lines show the global best-fit, with the corresponding residuals shown in the plot above. For clarity, every third experimental data point is displayed.

**Figure S2.**
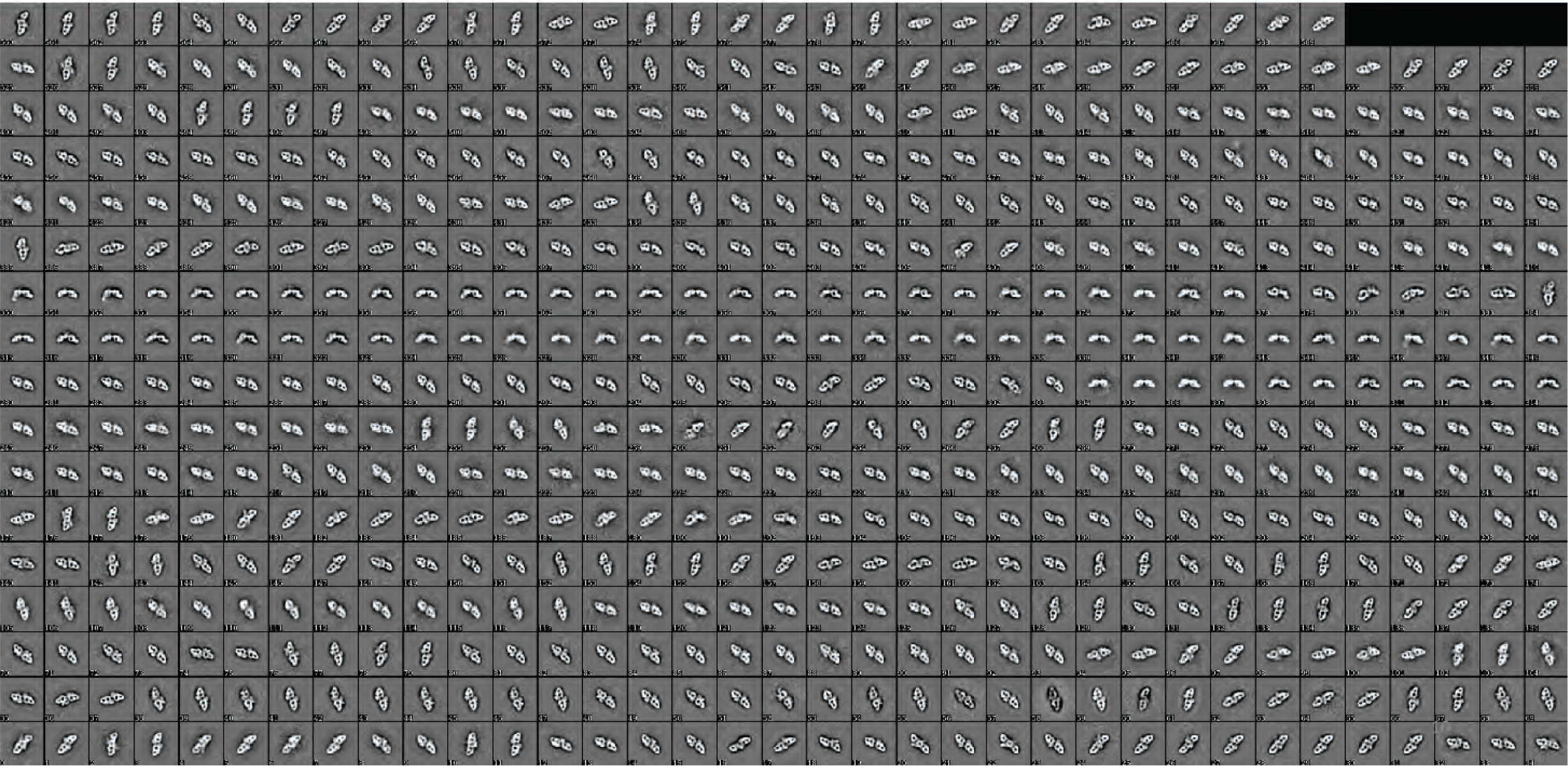
Back projection of Neurofibromin EM model to 2D class averages. The 3D EM model generated was used to calculate 2D back projections using EMAN2. A representative sample of the back projected 2D classes is shown here.

**Figure S3.**
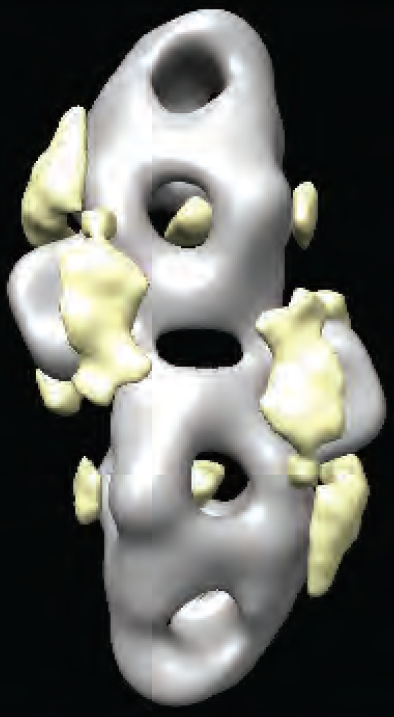
3D variability analysis of Neurofibromin structure. 3D variability analysis showing heterogeneity in the 3D model of full-length Neurofibromin. The heterogenous areas are shown in yellow.

**Figure S4.**
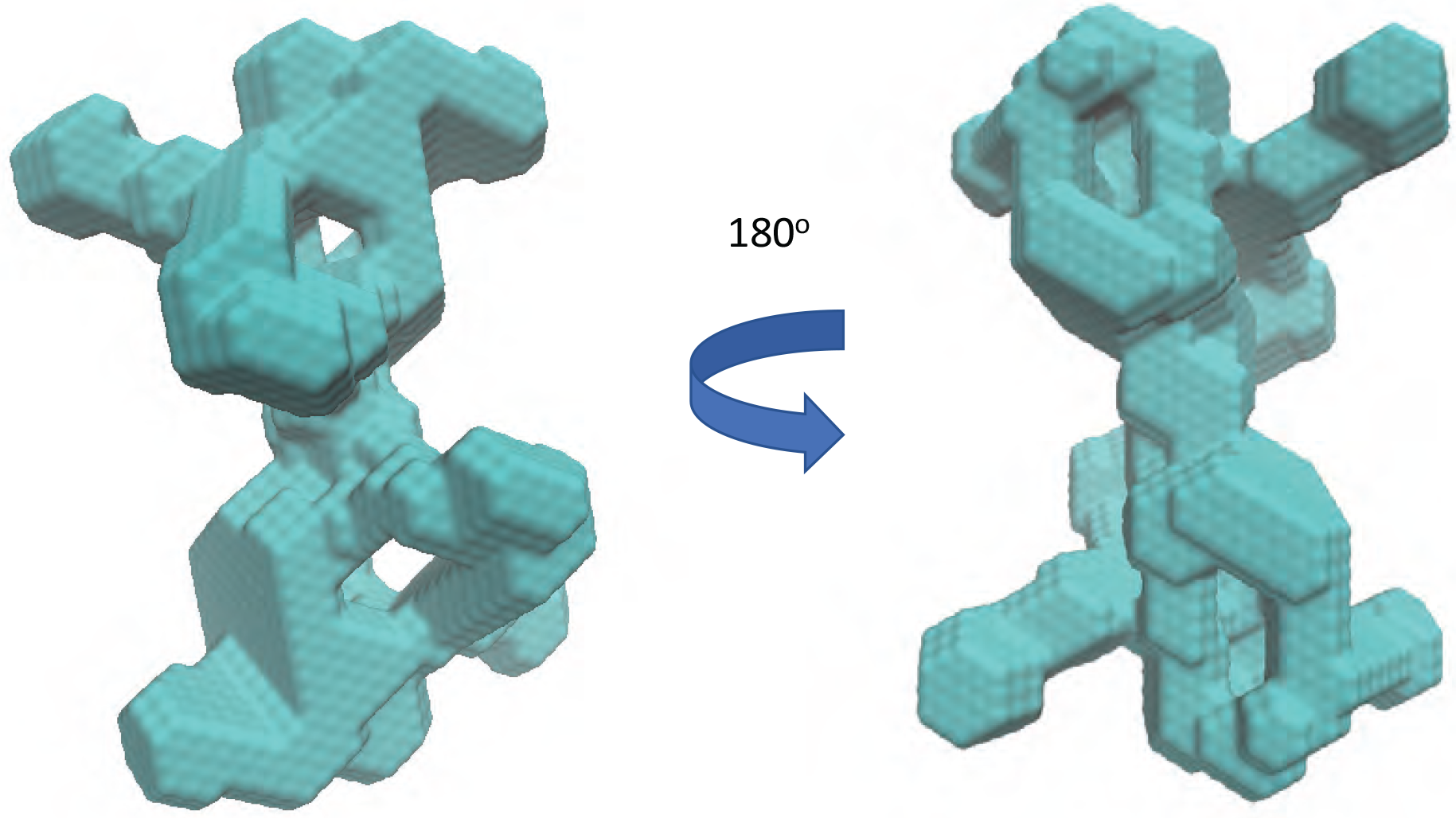
*Ab initio* model of Neurofibromin dimers using SAXS/SANS data. A representative NF1 *ab initio* shape reconstruction generated from the SAXS data.

**Figure S5.**
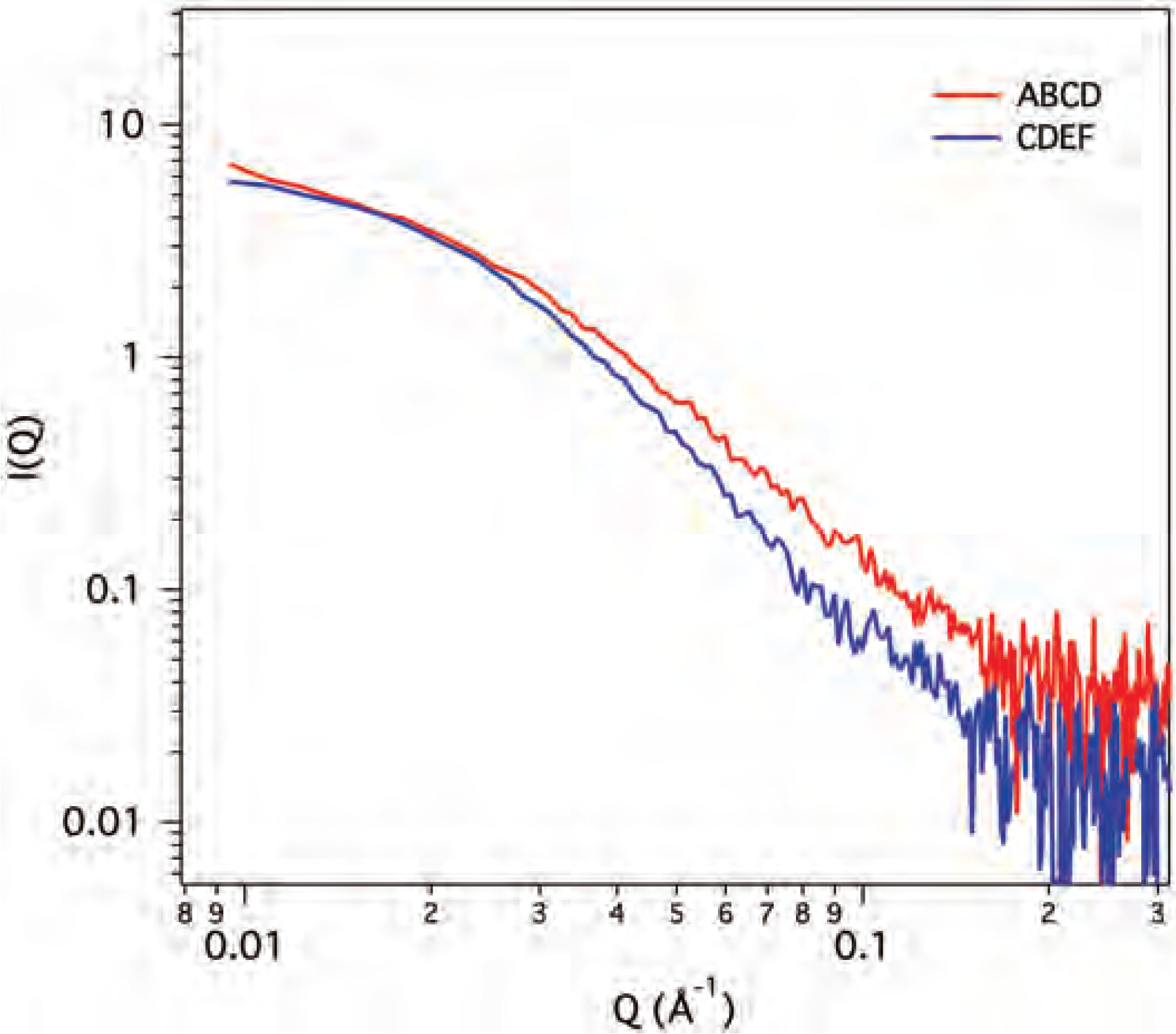
Small-angle X-ray scattering of ABCD and CDEF fragments of Neurofibromin. ABCD protein (red line) was measured at 1.5 mg/ml (7.5 uM) and CDEF protein (blue line) was measured at 4.3 mg/ml (16 uM).

**Figure S6.**
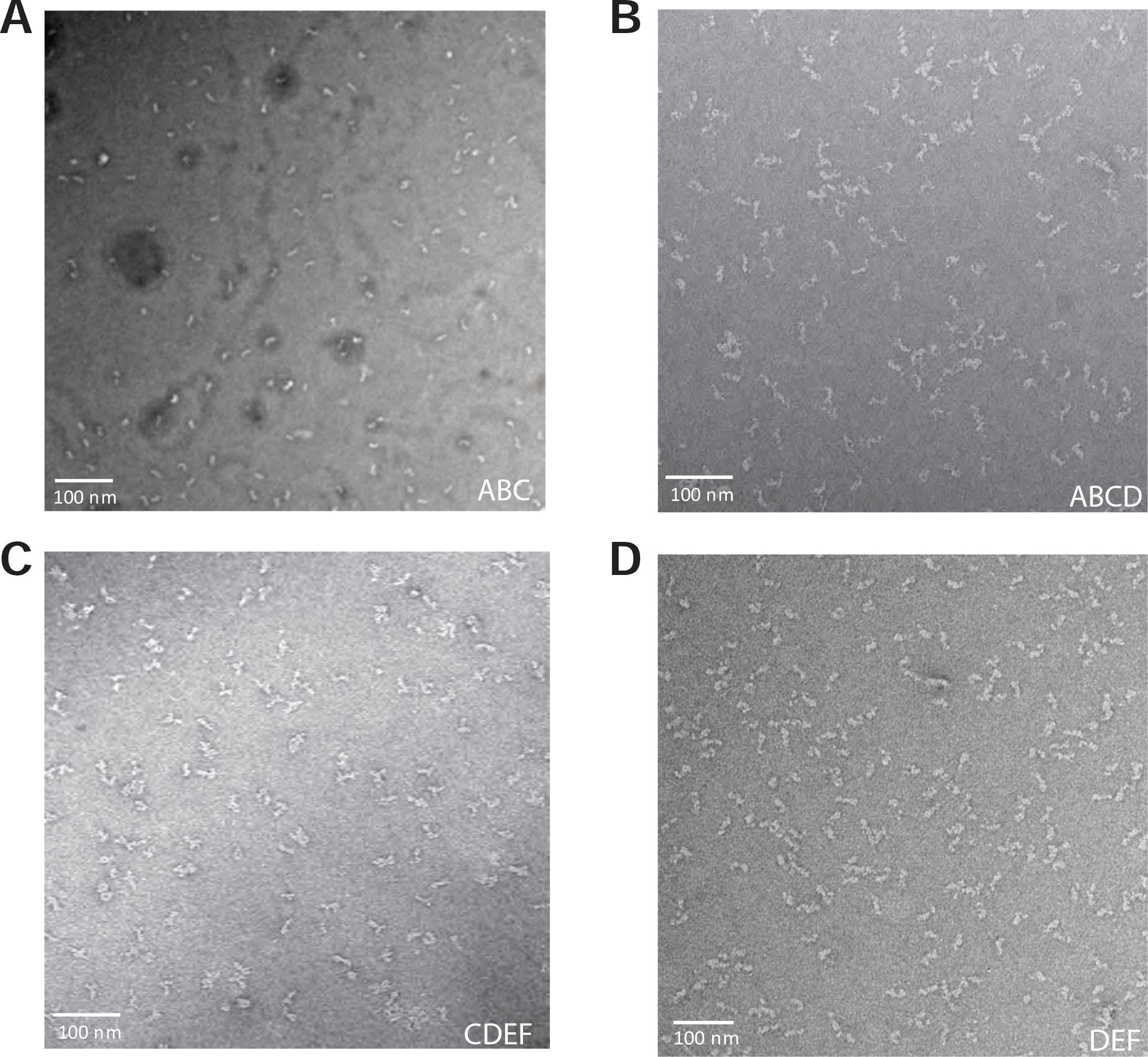
Negative stain transmission EM images of Neurofibromin domain proteins. Representative transmission electron micrographs of purified ABC (panel A), ABCD (panel B), CDEF (panel C), and DEF (panel D) proteins.

**Figure S7.**
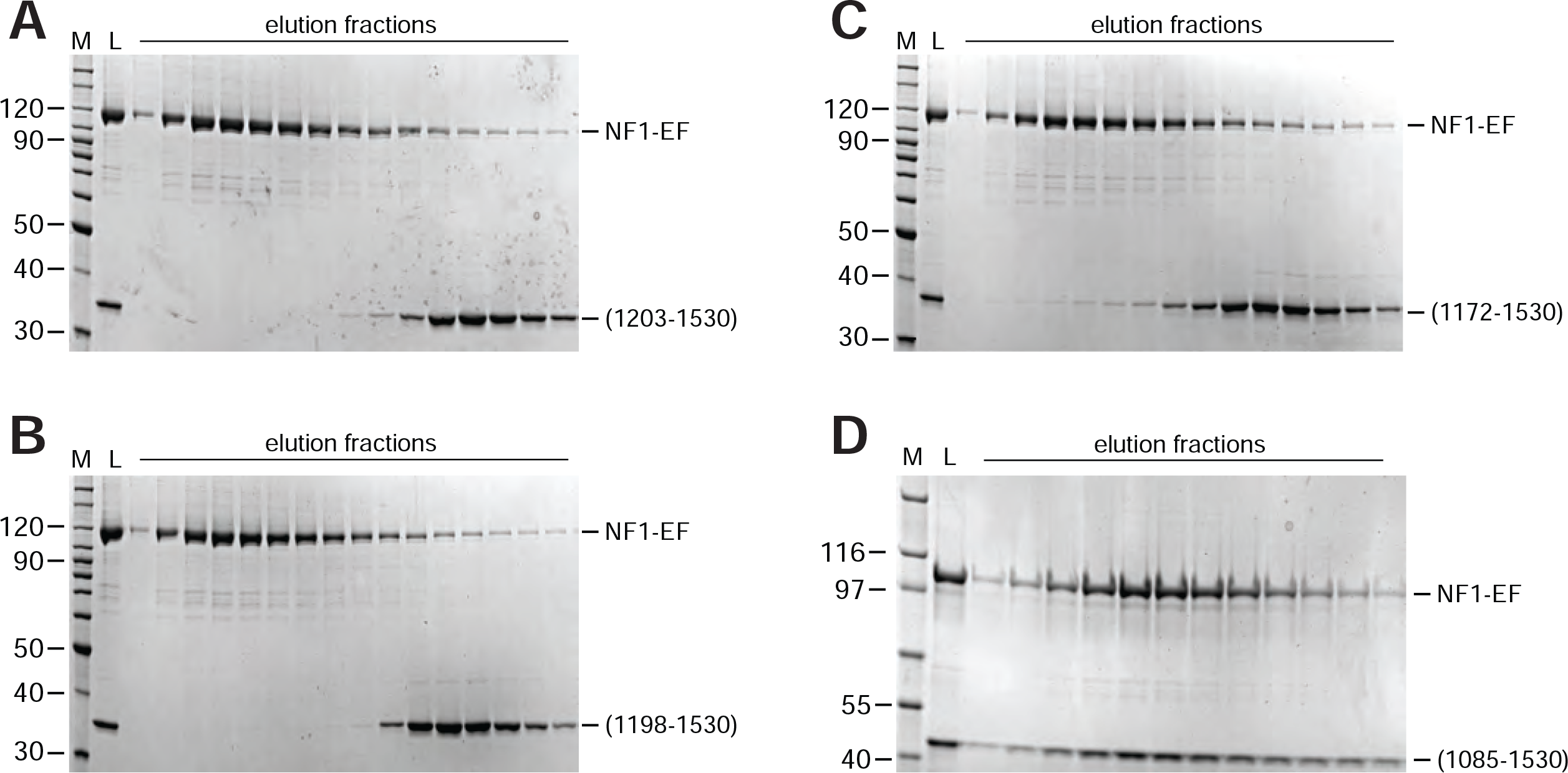
Formation of complexes of Neurofibromin C and EF domains requires the presence of the TBD region. SDS-PAGE analysis of SEC of equimolar mixtures of NF1 domains with EF (1821-2818). In each panel, the size of molecular weight markers in lane M is noted in kilodaltons. Lane L represents the loaded material, and additional lanes are elution fractions across the column. Neurofibromin fragments mixed with the EF domain in each panel were: *A*, 1203-1530; *B*, 1198-1530; *C*, 1172-1530; *D*, 1085-1530. Coeluting proteins are observed only in the construct containing amino acids 1085-1171.

**Figure S8.**
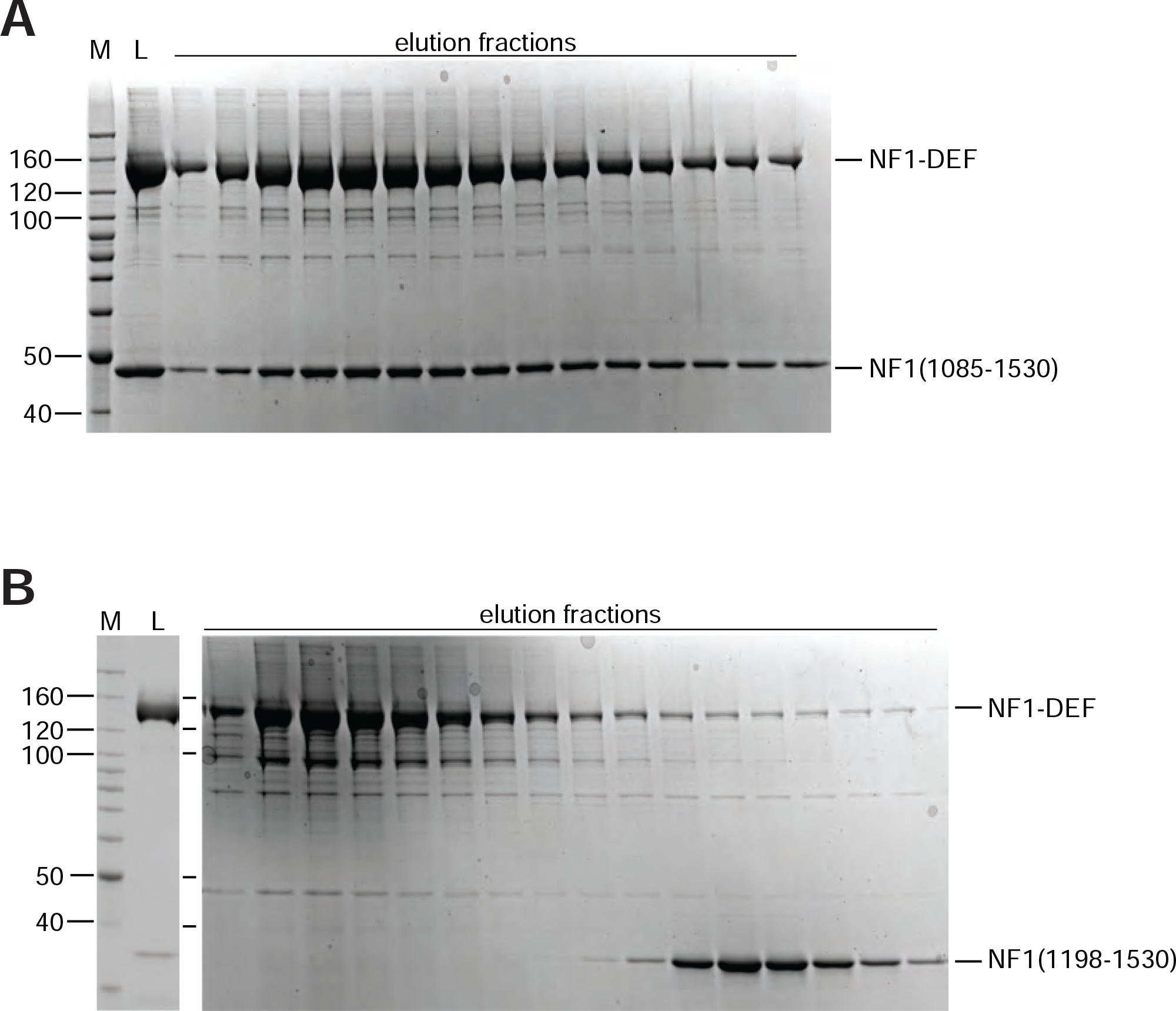
Formation of complexes of Neurofibromin C and DEF domains requires the presence of the TBD region. SDS-PAGE analysis of SEC of equimolar mixtures of NF1 domains with DEF (1541-2818). In each panel, the size of molecular weight markers in lane M is noted in kilodaltons. Lane L represents the loaded material, and additional lanes are elution fractions across the column. Neurofibromin fragments mixed with the DEF domain in each panel were: *A*, 1085-1530; *B*, 1198-1530. Coeluting proteins are observed only in the construct containing amino acids 1085-1197.

